# Integrative Transcriptomics and Phytochemical Screening Reveal Pratenol B, Eriodictyol, Losbanine, and Isookanin, as Potential EGFR and HRAS Inhibitors in Indian Oral Squamous Cell Carcinoma Patients

**DOI:** 10.64898/2025.11.28.691262

**Authors:** Shubha Behara, Uday Yadav, Vishal Kumar Sahu, Shuchi Nagar, Soumya Basu, Samir Gupta, B. M. Rudagi, Supriya Kheur, Dimple Davray, Subhayan Sur

**Affiliations:** Cancer and Translational Research Center, Dr. D. Y. Patil Biotechnology and Bioinformatics Institute, Dr. D. Y. Patil Vidyapeeth, Tathawade, Pune - 411033, India; Bioinformatics Centre, Dr. D. Y. Patil Biotechnology and Bioinformatics Institute, Dr. D. Y. Patil Vidyapeeth, Tathawade, Pune - 411033, India; Department of Surgical Oncology, Dr. D. Y. Patil Medical College, Hospital & Research Centre, Dr. D. Y. Patil Vidyapeeth (DPU), Pune-411018; Department of Oral & Maxillofacial Surgery & Oral Implantology, Dr. D. Y. Patil Dental College & Hospital, Dr. D. Y. Patil Vidyapeeth (DPU), Pune-411018; Department of Oral Pathology and Microbiology, Dr. D. Y. Patil Dental College & Hospital, Dr. D. Y. Patil Vidyapeeth (DPU), Pune-411018

**Keywords:** Oral Squamous Cell Carcinoma, Phytochemicals, Cancer therapy, EGFR, HRAS, Pratenol B

## Abstract

Oral squamous cell carcinoma (OSCC) is the most common head and neck cancer, with India contributing nearly one-third of the global cases. Management of OSCC remains difficult due to increasing risk factors, limited therapeutic options, severe side effects, and rising drug resistance. Therefore, novel and safer treatment strategies are urgently needed. This study explores the potential of phytochemicals as targeted inhibitors of key dysregulated biomarkers in Indian OSCC patients. RNA sequencing and pathway analysis revealed significant alterations in the MAPK signaling pathway, highlighting EGFR and HRAS as crucial therapeutic targets. Given the limited clinical success of existing EGFR-targeted therapies and the scarcity of HRAS inhibitors, a natural product-based approach was adopted. Molecular docking of 17,000 phytochemicals identified Pratenol B, Eriodictyol, Losbanine, and Isookanin as promising inhibitors, with Pratenol B showing dual inhibition of EGFR and HRAS. These compounds exhibited strong binding affinities, favorable pharmacokinetic profiles, high bioavailability, and low toxicity. Molecular dynamics simulations confirmed the stability of Pratenol B with both target proteins, surpassing reference inhibitors. Utilizing vast medicinal plant diversity presents a cost-effective and low-toxicity avenue for OSCC therapy. Further in vitro, in vivo, and clinical studies are warranted to validate these phytochemicals as potential therapeutics.

## 1. Introduction

Oral squamous cell carcinoma (OSCC) is the most prevalent head and neck cancer, affecting the tongue, gingivae, lips, cheeks, and oral cavity (Usman et al., 2020). Its incidence varies geographically and is primarily linked to tobacco use, excessive alcohol consumption, and betel nut chewing (Bray et al., 2024, Johnson et al., 2020). In South Central Asia, particularly India, OSCC accounts for nearly one-third of global cases due to high-risk behaviours (Borse et al., 2020, Bray et al., 2024). The Global Cancer Observatory (GCO, 2022) estimated 389,846 new cases of oral cancer, with a 48.33% mortality rate worldwide (Bray et al., 2024). In the United States, the estimated incidence for 2025 is 59,660 cases, with 12,770 deaths (Siegel et al., 2025). The annual increase in OSCC incidence by 1% and HPV-associated oral cancer mortality rising by 2% emphasize the need for early diagnosis and improved treatment options (Siegel et al., 2025).

OSCC is a multistage disease characterized by genetic alterations that drive abnormal proliferation, differentiation, survival, and metastasis. Key contributors include the overexpression of proto-oncogenes such as EGFR, HRAS, KRAS, NRAS, c-myc, and PRAD-1 (Usman et al., 2020). EGFR, often overexpressed in OSCC due to gene amplification, plays a critical role in tumor progression (Usman et al., 2020). The HRAS gene, a member of the Ras family, is frequently mutated in OSCC cases in Asian populations, especially among betel nut users (Usman et al., 2020). Current EGFR-targeting therapies (Cetuximab, Gefitinib, Erlotinib) show promise, but no FDA-approved treatments exist for HRAS mutations (Fasano et al., 2021, Guimond et al., 2022). Surgical resection and radiation therapy remain standard treatments, but they are associated with complications such as tumor seeding, organ dysfunction, and osteoradionecrosis. FDA-approved therapies like cetuximab, pembrolizumab, and nivolumab offer limited options, with side effects such as rashes, diarrhea, and immune-related reactions reducing their effectiveness (Fasano et al., 2021, Guimond et al., 2022, Naruse et al., 2016, Ferris et al., 2016, Harrington et al., 2023).

Plant-derived phytochemicals have been central to drug discovery, contributing to the development of many drugs including leucovorin, carzinophilin, and vincristine (Sur and Ray, 2023, Newman and Cragg, 2020). Nearly 50% of anticancer drugs originate from natural sources, including vinca alkaloids, taxanes, and anthracyclines (Safarzadeh et al., 2014). Between 1981 and 2019, an approximately 1,881 of FDA-approved drugs were derived from natural products (Newman and Cragg, 2020). Plant secondary metabolites offer cost-effective, low-toxicity alternatives to synthetic drugs, improving cancer therapy through enhanced bioavailability and tumor specificity. Phytochemicals, either alone or in combination with conventional treatments, have shown promise in reducing tumor growth, inhibiting cell proliferation, and alleviating cancer-related pain (Burcher et al., 2023).

Thus, this study aimed to identify potential phytochemicals targeting key OSCC biomarkers. RNA sequencing of patient samples revealed that the MAPK signaling pathway was the most significantly altered in tumors. Based on fold changes and pathway interactions, EGFR and HRAS were prioritized as therapeutic targets. Molecular docking and binding energy calculations identified potential phytochemicals for these targets, which were further assessed for pharmacokinetics, bioactivity, and bioavailability to determine their drug-likeness. The findings highlight promising plant-derived inhibitors for EGFR and HRAS, offering a potential breakthrough in OSCC treatment with fewer side effects and greater accessibility.

## 2. Materials and Methods

### 2.1. OSCC Patient Sample Collection

OSCC patient samples, along with adjacent non-tumor tissues, were collected with informed consent from Dr. D.Y. Patil Dental College & Hospital and Dr. D.Y. Patil Medical College, Hospital & Research Centre, DPU, Pune. The study was approved by the Ethics Committee of Dr. D.Y. Patil Vidyapeeth (DPU), Pune and Institutional Biosafety committee.

### 2.2. Analysis of Transcriptomic Profile in OSCC Patient Samples

To examine the global transcriptomic profile, RNA sequencing was performed on three tumor samples (T1, T2, and T3) and three adjacent non-tumor samples (N1, N2, and N3), as previously described (Sur et al., 2022). Total RNA was isolated using TRIzol reagent (Invitrogen), and cDNA was synthesized using the iScript cDNA synthesis kit (BioRad). Sequencing was conducted on the Illumina platform, with alignment performed using Hisat2 (version 2.1.0) tools. Gene abundance was estimated with FeatureCounts (2.0.1). Gene IDs for both non-coding and protein-coding genes were categorized using Ensembl Biomarts. Differential expression analysis was carried out using R/Bioconductor package: DESeq2, by using negative binomial distribution based variance-mean dependence model to handle biological and technical variability. A p-value < 0.05 was considered significant, with upregulated genes defined by a log2 fold change ≥ 1 and downregulated genes by a log2 fold change ≤ -1.

### 2.3. Gene Ontology and Pathway Analysis

To classify significantly differentially expressed genes (DEGs) into predefined functional categories (Biological Process (BP), Cellular Component (CC), and Molecular Function (MF), Gene Ontology (GO) enrichment analysis was performed using clusterProfiler and topGo of the R/Bioconductor package (Sur et al., 2022). Further to pin down over represented biological pathways associated with differentially expressed genes (DEGs), KEGG pathway enrichment analysis was conducted to gain the functional insights, as outlined in previous work (Sur et al., 2022).

### 2.4. Target Selection and Preparation

The receptors are the biological target on which the ligands bind. In this study, receptors were selected from hub genes having high protein-protein interaction (PPI) score with highest log fold change among significantly differential genes expression. A high interaction score indicates string connectivity of receptor gene with other genes, suggesting its role as a key regulatory receptor. The selected 3D crystal structures of EGFR (PDB ID: 8A27) and HRAS (PDB ID: 8TBG) were obtained from the Protein Data Bank (PDB) based on their high resolution and taxonomic relevance (Obst-Sander et al., 2022, Holderfield et al., 2024). Before docking, the structures were refined by removing water molecules, unessential chains, and heteroatoms (Burley et al., 2017). Active binding sites were defined based on known active site residues and the presence of co-crystal ligands.

### 2.5. Selection and preparation of Phytochemical Ligands

In this study, phytochemicals were used as ligands due to their potential to bind with the selected biological targets. Till June 2024, the 3D structure of all available phytochemicals was obtained in SDF format from the IMPPAT (Indian Medicinal Plants, Phytochemistry, and Therapeutics) database, chosen for its comprehensive coverage of traditional Indian medicinal plants (Mohanraj et al., 2018, Vivek-Ananth et al., 2023). To evaluate the drug-like properties, these compounds were subjected to a screening process and ranked them from 1 to 5 based on Lipinski’s Rule of Five. The screening and ranking process were performed by using in-house Python scrips, available at the following repository: GitHub Link https://github.com/udayshivajiyadav/Phytochemicals-Screening-/tree/main.

### 2.6. Molecular Docking Simulations

Molecular docking simulations were performed using FlexX (LeadIT suite, BioSolveIT GmbH), a fragment-based incremental construction algorithm that docks ligands into defined active sites of target proteins (Pagadala et al., 2017, Lokhande et al., 2022). This approach enables predicting ligand orientation and interaction modes, maintaining receptor rigidity while allowing ligand flexibility. Docking scores were ranked, and top-ranking poses were visually inspected for key interactions such as hydrogen bonding, π-π stacking, and hydrophobic contacts. In cases of multiple docking poses, cluster analysis was used to determine the most probable binding mode.

### 2.7. In Silico ADME and Toxicity Predictions

Pharmacokinetic parameters, including ADMET (Absorption, Distribution, Metabolism, Excretion, and Toxicity) properties were predicted using the ADMETLab 3.0 API (Xiong et al., 2021). Toxicity predictions, including AMES toxicity, hepatotoxicity, and hERG inhibition, were conducted using pkCSM to evaluate potential mutagenic, liver-toxic, or cardiotoxic effects.

### 2.8. Bioactivity and Bioavailability Assessment

The bioactivity score of each ligand was determined using Molinspiration software, assessing interactions with GPCRs, enzyme inhibitors (EI), kinase inhibitors (KI), nuclear receptor ligands (NRL), and ion channel modulators (ICM) (Mendie and Hemalatha, 2022). Additionally, SwissADME was used to generate bioavailability radar plots, instantly evaluating oral bioavailability by analysing lipophilicity, polarity, molecular size, solubility, flexibility, and unsaturation.

### 2.9. Molecular Dynamics Simulation

To evaluate the interaction and stability of EGFR and HRAS with the top three ligands (selected based on drug-likeness and bioavailability properties), a 100-nanosecond molecular dynamics (MD) simulation was performed using the Desmond module of Schrödinger software. Molecular dynamics simulation enables the calculation of forces and the motion of amino acid atoms, offering insights into receptor-ligand interactions. Desmond integrates parameters such as system volume, pressure, and temperature, and is equipped with features specifically designed for detailed receptor-ligand interaction studies.

Energy minimization, a critical step in MD simulations, was carried out using the steepest descent method. The solvated complex system was neutralized by adding appropriate counterions (Na□ and Cl□). To minimize edge effects while applying periodic boundary conditions, an orthorhombic simulation box with a buffer size of 10 Å was used. The TIP3P water model, in combination with the OPLS-AA force field—suitable for enzyme simulations—was employed to model the solvent environment and calculate the system’s potential energy (Jorgensen et al., 1996).

System equilibration was achieved under constant temperature (300 K) and pressure (1.01325 bar) using the Nosé–Hoover thermostat and Martyna–Tobias–Klein barostat. After equilibration, the simulation trajectories were collected and analyzed to assess the interaction stability of the receptor-ligand complexes (Evans and Holian, 1985, Janek and Kolafa, 2024).

Post-simulation analysis was conducted using Desmond’s Simulation Interaction Diagram (SID) and Simulation Event Analysis (SEA) tools. Key parameters such as root mean square deviation (RMSD) and root mean square fluctuation (RMSF) were calculated from the 100 ns trajectory data. These analyses provided insights into the conformational stability and fluctuations of the receptor-ligand complexes, by comparing the initial docked structures with their dynamic behavior over the simulation time.

## 3. Results and Discussion

### 3.1. Global Transcriptomic changes in OSCC patient samples

The prevalence, mortality, and molecular mechanisms of OSCC can vary significantly across geographical regions, influenced by factors such as prevalent risk behaviours, environmental exposures, and genetic predispositions (Ram et al., 2011, Gormley et al., 2022). In India, OSCC is highly prevalent, with genetic and environmental factors contributing to differences in its occurrence, diagnosis, and treatment compared to the Western world (Borse et al., 2020).

To better understand the molecular mechanisms, RNA sequencing analysis was performed to examine the global transcriptomic profile in OSCC samples. Both OSCC and non-tumor samples were validated histopathologically (data not shown). The samples were collected from male individuals with an average age of 47 years, all of whom had a history of tobacco use, tested negative for viral infections, and were diagnosed with well-differentiated carcinoma. The key metrics for the sequencing are presented in **Supplementary Figure 1A and 1B**. Our RNA sequencing data generated a total of 40,587 mRNA genes, of which 26,597 genes were significantly modulated in OSCC samples compared to non-tumor tissue (**Figure1 A, B, and Supplementary Figure 1C**). Among these, 7,738 genes were significantly upregulated, and 13,505 were significantly downregulated (**Supplementary Figure 1C**). The gene expression of some of the most significantly modulated genes is shown in **Figure 1C**.

**Figure 1:**
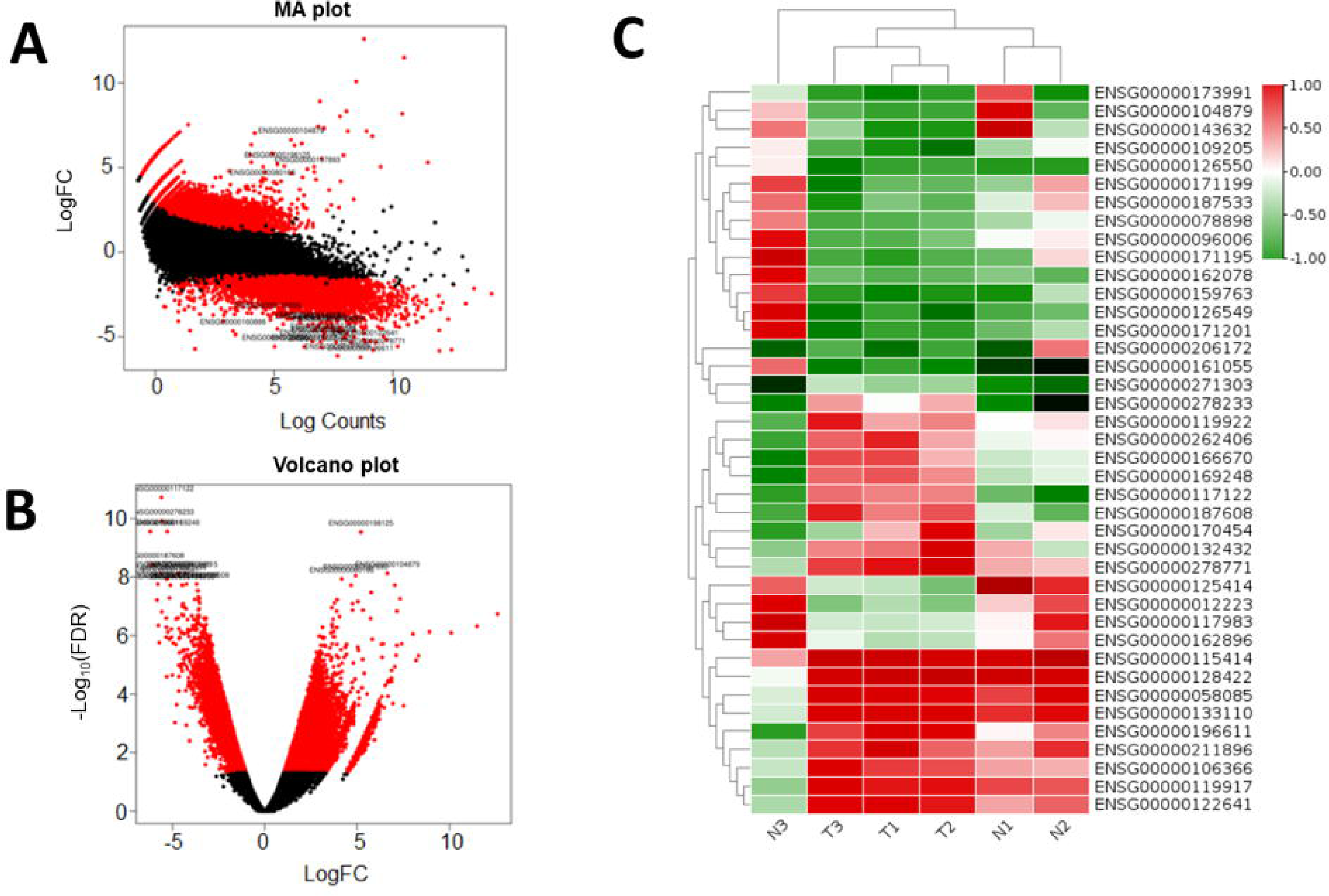
Global Transcriptomic Profile of Indian OSCC Patient Samples. (A) MA plot illustrating global gene expression changes, where red dots represent significantly differentially expressed genes. **(B)** Volcano plot highlighting significantly upregulated and downregulated genes (red dots), providing insights into the most prominent transcriptomic alterations in OSCC. **(C)** Heatmap depicting sample-wise (Tumor samples: T1-3 and adjacent non-tumor samples: N1-3) expression variations of the top differentially expressed genes, showcasing distinct molecular signatures in OSCC patient samples.

### 3.2. Gene Ontology and pathway analysis

Next, Gene Ontology (GO) and Kyoto Encyclopedia of Genes and Genomes (KEGG) pathway analysis was performed in OSCC using the R/Bioconductor package. **Figure 2A-C** present the GO analysis under the categories of ‘Biological Process’, ‘Cellular Component’, and ‘Molecular Function’. In the ‘Biological Process’ category, key processes such as ‘Biological Regulation’, ‘Metabolic Process’, ‘Cell Communication’, and ‘Growth’ were identified. For ‘Cellular Component’, the most significantly altered components were associated with the ‘Membrane’ and ‘Nucleus’. In terms of ‘Molecular Function’, the predominant functions were ‘Protein Binding’, ‘Ion Binding’, and ‘Nucleic Acid Binding’.

**Figure 2:**
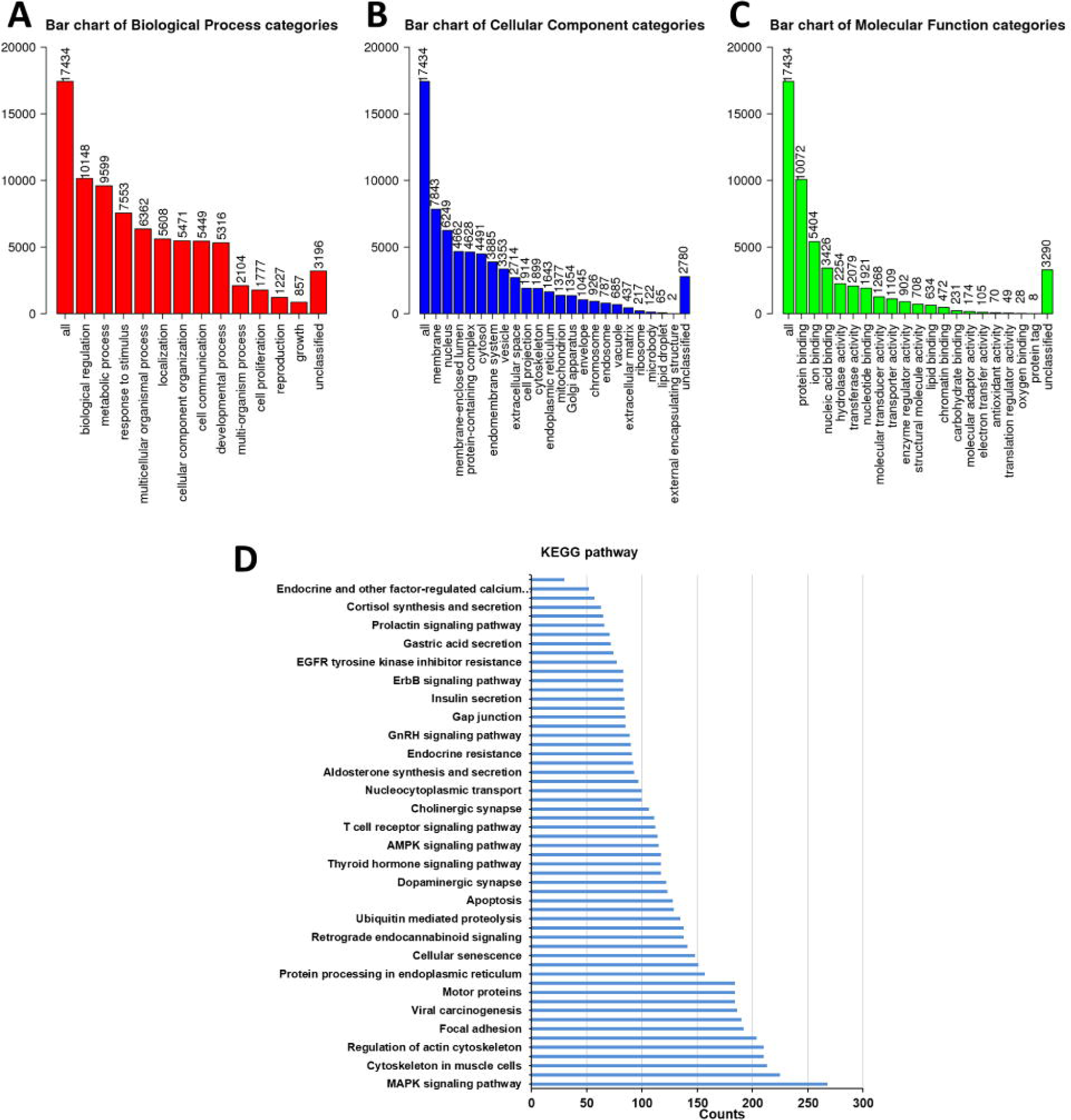
Alterations in Gene Ontology Categories and KEGG Pathways in OSCC Patient Samples. (A–C) Gene Ontology (GO) enrichment analysis highlighting significantly modulated categories across Biological Process (BP), Cellular Component (CC), and Molecular Function (MF) in OSCC patient samples. **(D)** KEGG pathway analysis identifying significantly dysregulated pathways, with MAPK signaling emerging as the most prominently altered pathway, suggesting its critical role in OSCC pathophysiology.

KEGG pathway analysis identified the MAPK signaling pathway (hsa04010) as the most significantly altered pathway in OSCC samples (**Figure 2D**). Other significantly altered pathways are: ‘hsa04110: Cell cycle’, ‘hsa04935: Growth hormone synthesis, secretion and action’, ‘hsa04152: AMPK signaling pathway’, ‘hsa04066: HIF-1 signaling pathway’, ‘hsa04012: ErbB signaling pathway’, ‘hsa04514: Cell adhesion molecules’, ‘hsa01521: EGFR tyrosine kinase inhibitor resistance’, ‘hsa04115: p53 signaling pathway’, ‘hsa04370: VEGF signaling pathway’; ‘hsa00020: Citrate cycle (TCA cycle)’ etc. (**Figure 2D**).

The mitogen-activated protein kinase (MAPK) signaling pathway is involved in tumor cell proliferation, differentiation, apoptosis, angiogenesis, invasion, metastasis and associated with patient prognosis (Peng et al., 2018, Cheng et al., 2022). Activation of the MAPK signaling pathway occurs in over 50% of OSCC cases, with mutations being more prevalent in Asian populations (Cheng et al., 2022, Peng et al., 2018). Thus, a thorough understanding of MAPK signaling may be crucial for elucidating the molecular mechanisms underlying the pathogenesis of OSCC in this population.

### 3.3. Identification of potential targets

The MAPK signaling pathway was closely examined to understand the molecular basis of oral cancer. The significantly modulated genes in this pathway, including VEGFC, CDC25B, AREG, EGFR, and HRAS, are shown in **Figure 3A**. The clinical significance of the top ten significantly modulated genes, along with their expression changes, molecular mechanisms, and available drugs, is presented in **Table 1**. The functional interactions of the top 20 genes in the pathway are depicted in **Figure 3B**, analyzed using STRING analysis. This indicates importance of MAPK signalling molecule in OSCC of Indian patient.

**Figure 3:**
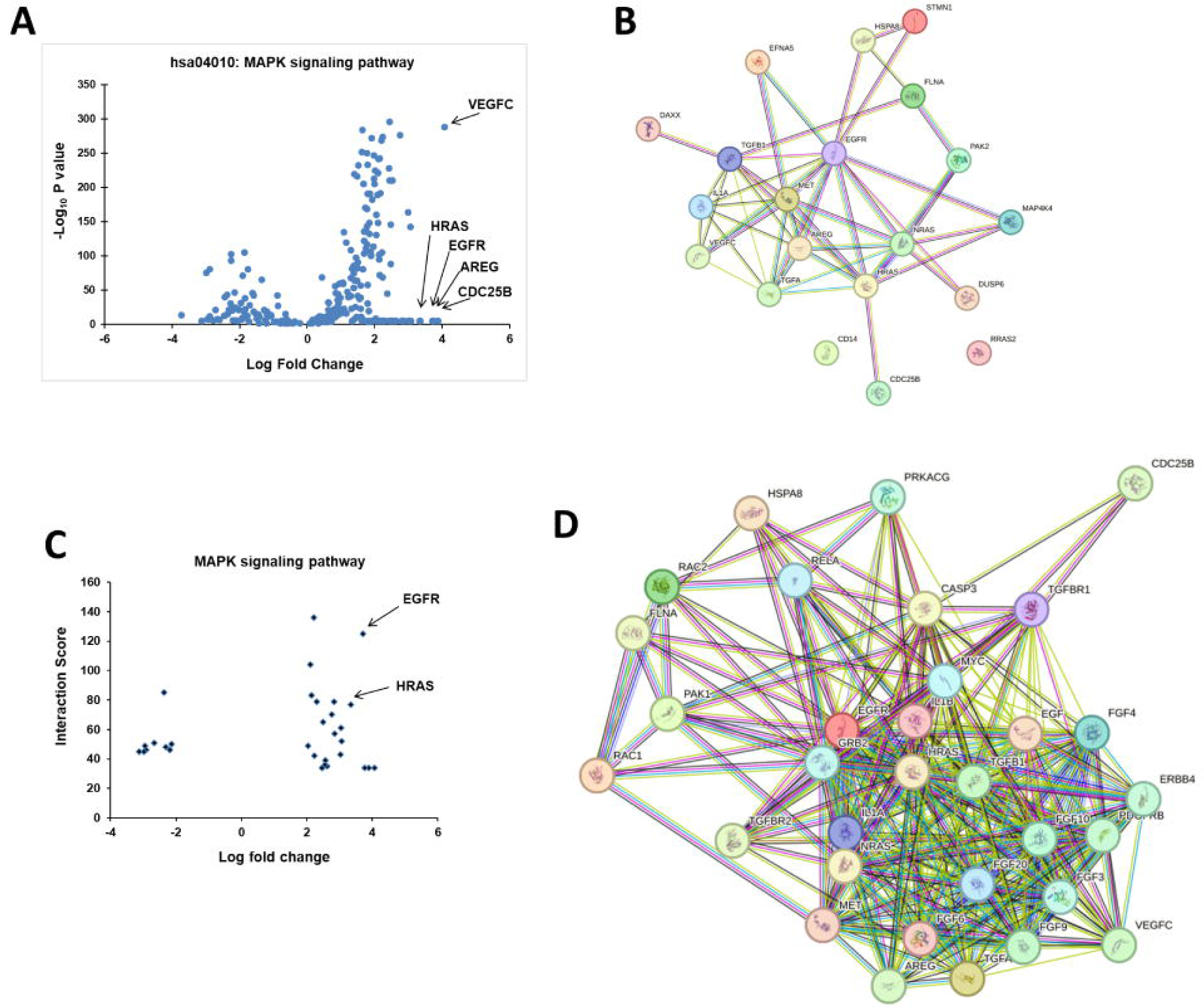
Dysregulation and Functional Network Analysis of the MAPK Signaling Pathway in OSCC. **(A)** Volcano plot illustrating differential gene expression within the MAPK signaling pathway, with arrows highlighting key upregulated genes. **(B)** STRING network analysis depicting functional interactions among the top 20 differentially expressed genes, as identified using STRING database analysis. **(C)** Volcano plot representing hub genes identified through STRING-based network analysis, ranked based on interaction scores and fold-change expression in OSCC samples. **(D)** Functional interaction network of hub genes within the MAPK signaling pathway, emphasizing key molecular interactions and their potential role in OSCC pathogenesis.

**Table 1:**
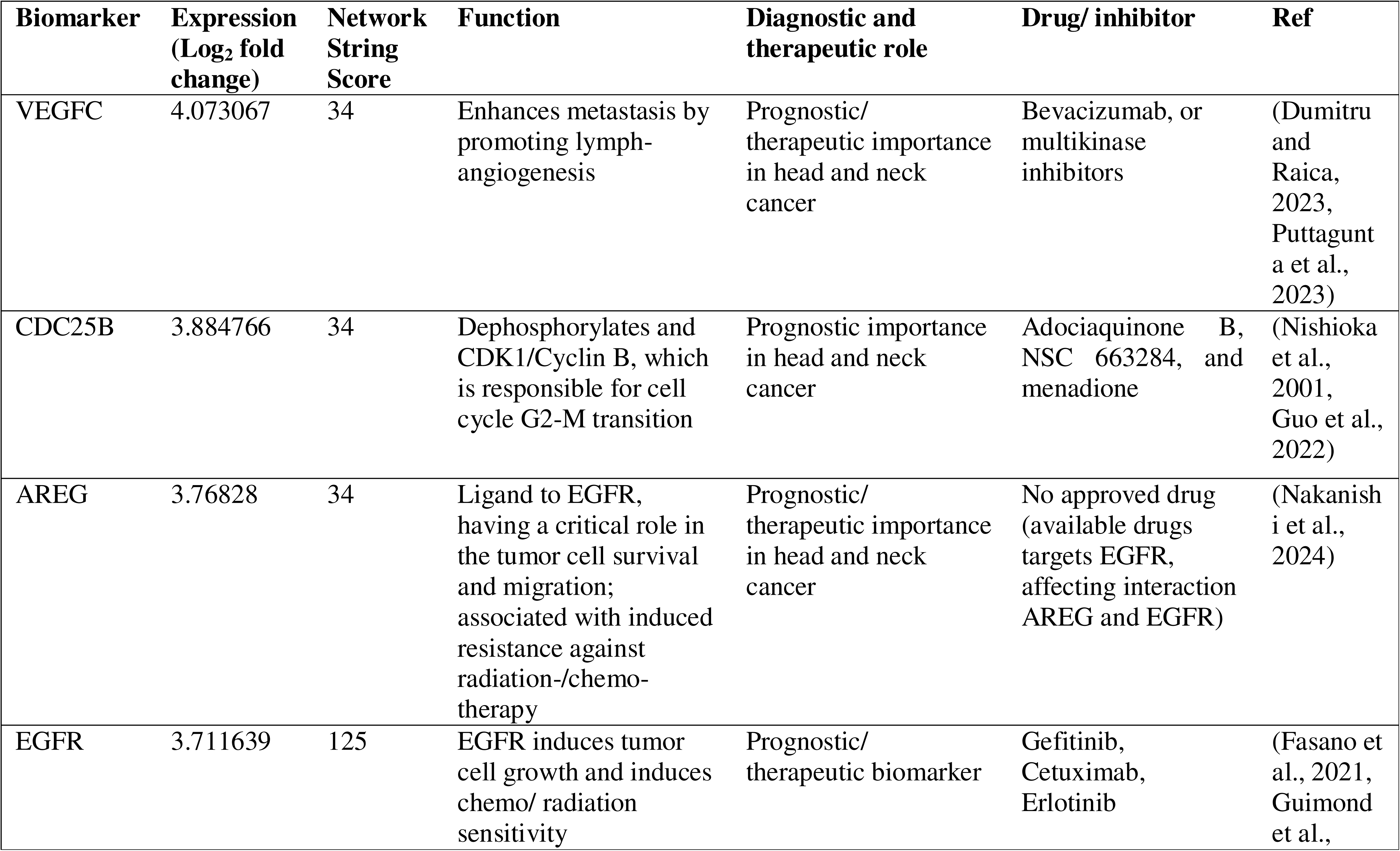
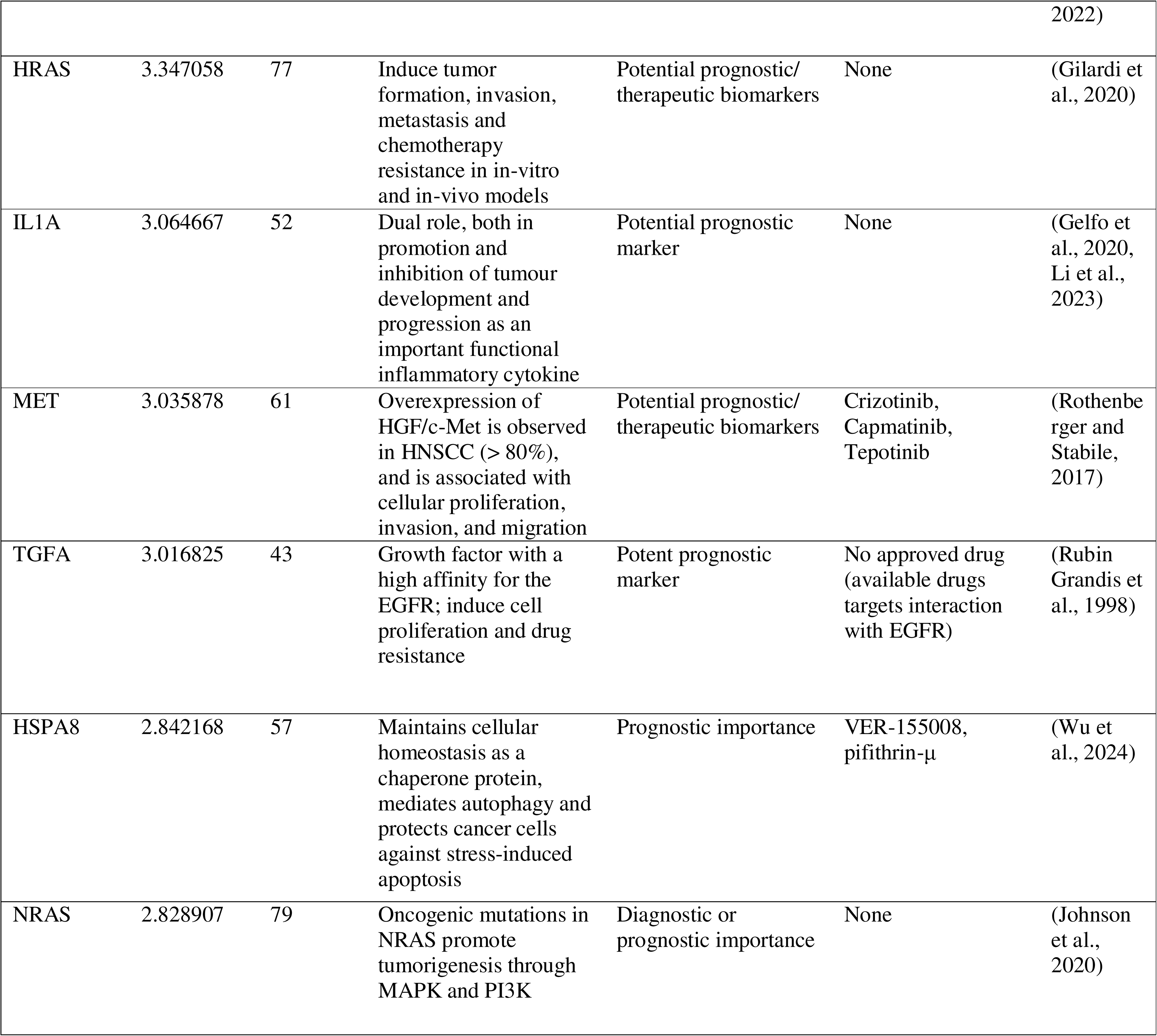
Importance of some top significantly expressed genes from MAPK signaling pathway in head and neck cancer or OSCC

The MAPK signaling molecules were also examined to create a hub gene network, mapping the interactions and relationships between genes and proteins in the given biological context. This approach highlights key players (hub genes) that may offer insights into disease mechanisms and potentially guide therapeutic strategies. The genes, along with their interaction scores, are shown in **Figure 3C** and **Table 1**, while their STRING functional associations are presented in **Figure 3D**. Based on the top interaction scores and fold changes, we selected EGFR and HRAS as the top two molecules for further analysis.

The role of EGFR and HRAS in oral cancer is highlighted in **Table 1**. Both molecules are upregulated in OSCC, promoting cell proliferation and contributing to drug resistance. Studies have suggested their prognostic and therapeutic significance.

As discussed above, genetic variation among human populations is influenced by geographical distribution, where the Indian population exhibiting distinct genomic differences compared to other global populations (Borse et al., 2020, Xing et al., 2010). These population-specific genetic distinctions highlight the relevance of precision medicine, which focuses on identifying cancer-specific genetic alterations to enable personalized therapeutic interventions.

In the Indian population, overexpression, frequent amplification, and prognostic significance of the EGFR gene have been well documented (Mohanapure et al., 2022, Gore et al., 2011, Singla et al., 2018). Additionally, mutations in the HRAS gene, which are commonly observed in OSCC cases across Asian populations, have also been reported in Indian OSCC patients (Usman et al., 2020, Muthusamy et al., 2025, Muthusamy et al., 2023). These reports are consistent with our own observations in patient samples, suggesting importance of EGFR and HRAS in the pathogenesis of OSCC within this population.

While therapeutic inhibitors and monoclonal antibodies targeting EGFR are available, no approved drugs currently target HRAS (**Table 1**). The use of EGFR inhibitors, such as cetuximab, in head and neck or OSCC is limited by challenges such as tumor heterogeneity, resistance, and significant side effects like skin toxicity and gastrointestinal symptoms (Sharafinski et al., 2010). Additionally, the lack of reliable predictive biomarkers and the high cost of treatment hinder their widespread efficacy and accessibility (Sharafinski et al., 2010). On the other hand, although mutations or overexpression of HRAS are common, and HRAS could serve as a valuable prognostic or predictive biomarker, no drugs are currently available to target HRAS. Therefore, it is crucial to identify potential small molecule inhibitors with fewer side effects for EGFR and HRAS.

### 3.4. Target selection

India is a land rich in diverse medicinal plants, many of which have been used in Ayurveda since ancient times (Pandey et al., 2013). It is now well-documented that a significant number of modern drugs are derived from plant-based phytochemicals (Chaachouay and Zidane, 2024). The advantages of these natural compounds include cost-effectiveness and minimal side effects (Chaachouay and Zidane, 2024). Therefore, we aimed to explore natural resources as a potential solution to address the challenges in targeting EGFR and HRAS.

The 3D crystal structures of human EGFR and HRAS proteins were retrieved from the Protein Data Bank (PDB), based on their recent upload dates, taxonomic relevance, and high resolution. The resolutions of these structures are 1.07 Å and 1.20 Å, respectively. The protein structures were refined and processed for docking analysis by removing water molecules, redundant chains, and heteroatoms (Burley et al., 2017).

### 3.5. Docking Analysis

Binding affinity between ligands and receptors is determined by binding energy, where lower energy indicates higher affinity (Mendie and Hemalatha, 2022). Of the 17,000 molecules, 10,763 phytochemicals from IMPPAT met all five of Lipinski’s rules, making them ideal drug-like candidates. Molecular docking simulations were performed using the FlexX module from the LeadIT suite (BioSolveIT GmbH). FlexX evaluates ligand-protein interactions such as hydrogen bonds, van der Waals forces, and electrostatic interactions to predict binding affinity. The docking poses were ranked based on their scores, and the top poses were further analyzed for key interactions, including hydrogen bonding, π-π stacking, and hydrophobic contacts.

The binding energy scores and interaction patterns were compared across ligands and reference ligands to identify potential high-affinity binders, making this method a valuable tool for rational drug design. The top 10 phytochemicals, selected based on their binding energy scores for the target EGFR and HRAS, are listed in **Table 2** along with their IDs and sources. Their chemical structures are shown in **Supplementary Figure 2**. Among these, Pratenol B (IMPHY009295), a benzopyran organo-heterocyclic compound, is common to both EGFR and HRAS, with binding energy scores of -24.7 and -50.6 kcal/mol, respectively. It is derived from the aerial parts of the Fabaceae family herb, *Trifolium pratense* (red clover) (**Table 2**). Additionally, (-)-Eriodictyol (R), which binds to EGFR, and Eriodictyol (S), which binds to HRAS, are enantiomers (**Table 2**). Interestingly, the binding energies of the ligands for EGFR and HRAS are comparable to those of the reference molecules—Gefitinib and the co-crystallized ligand phosphoaminophosphonic acid-guanylate ester, respectively (**Table 2**). Notably, the binding energies of the top 10 phytochemical ligands for EGFR are higher than those of the reported EGFR inhibitors, including Gefitinib and Erlotinib, further supporting our observations (Kamal et al., 2023).

**Table 2:** Binding parameters between ligands and target proteins

| Name of ligands | Source | IMPPAT ID | Binding Energy (Kcal.Mol) | No. of Hydrogen Bonds | Amino acids involved in Hydrogen Bonds |
| --- | --- | --- | --- | --- | --- |
| Target protein EGFR |  |  |  |  |  |
| Moschamine | *Carthamus tinctorius* (Safflower) -seed | IMPHY006312 | -29.199 | 7 | THR 854, ASP 855, PHE 856, GLY 857, LEU 858 |
| (-)-Eriodictyol (R)<br>Δ* | *Pistacia chinensis* (Chinese pistache)-fruit | IMPHY003897 | -29.042 | 12 | LEU 777, THR 790, THR 854, ASP 855, PHE 856, GLY 857, ALA 743, MET 766, CYS 775 |
| cis-Zeatin | *Ipomoea batatas* (Sweet potato) | IMPHY003896 | -27.569 | 6 | THR 854, ASP 855, LEU 788 |
| Anthragallol | *Hymenodictyon orixense* (Bridal couch tree)- roots | IMPHY001935 | -25.989 | 8 | LEU 777, THR 854, ASP 855, CYS 775, MET 766 |
| Robustone | *Derris robusta* (Beech Tree)- root and seed, and *Erythrina variegata* | IMPHY009228 | -25.856 | 4 | THR 854, ASP 855, GLY 857, CYS 775 |
| Adifoline | *Haldina cordifolia* (Haldu)-woods | IMPHY000977 | -25.726 | 8 | LYS 745, GLY 857, ILE 759, ASP 855, THR 854 |
| Vulgarol | *Artemisia indica*-rhizome, *Nothapodytes nimmoniana*- whole plant, *Tanacetum dolichophyllum*, and *Tarchonanthus camphorates*- leaf | IMPHY008954 | -25.645 | 5 | LYS 745, ASP 855 GLY 857, THR 854 |
| Losbanine * | *Alstonia scholaris* (Blackboard tree) – bark and leaf | IMPHY008426 | -25.629 | 5 | LYS 745, THR 854, ASP 855, THR 790 |
| 2-Anilino-4-phenylimino-naphthalen-1-one | *Reseda luteola* (Dyer's weed)-aerial part | IMPHY008417 | -25.294 | 1 | ASP855 |
| Pratenol B<sup>#</sup>* | *Trifolium pretense* (aerial part) | IMPHY009295 | -24.7529 | 5 | THR 854, ASP 855, PHE 856, ILE 759, LYS 745 |
| Gefitinib (Reference Molecule) | -- | -- | -20.732 | 6 | THR 854, ASP 855, ILE 759, GLY 857 |
| Target protein HRAS |  |  |  |  |  |
| Isookanin* | *Acacia leucophloea*-wood, and *Albizia julibrissin* | IMPHY009403 | -52.15 | 14 | GLY 13, GLY 15, GLY 60, VAL 14, GLN 61, ASP 33, THR 35, GLU 31 |
| Pratenol B<sup>#</sup> * | *Trifolium pratense* (Red clover)- aerial part | IMPHY009295 | -50.576 | 10 | GLY 60, THR 35, GLY 13, GLY 15, VAL 14 |
| Cedeodarin | *Cedrus deodara* (Deodar cedar)- flower and wood | IMPHY003355 | -48.924 | 10 | GLY 15, GLY 60, GLN 61, SER 17, LYS 16, THR 35, GLU 31, ASP 33 |
| Diferulic acid | *Prunus avium* and *Prunus cerasus*-shoot | IMPHY005552 | -48.075 | 11 | ASP 33, GLU 31, SER 145, ALA 146, LYS 147, GLY 60, THR 35, GLY 13 |
| 6-Amino-4-hydroxy-2-naphthalenesulfonic acid | *Humulus lupulus* (Common hop)- whole plant | IMPHY013746 | -46.75 | 13 | SER 17, VAL 14, LYS 16, GLY 13, GLY 15, ASP 33, GLU 31, VAL 29, ASP 30 |
| Padmatin | *Dittrichia graveolens*- aerial part, *Prunus cerasoides*- wood | IMPHY009332 | -46.094 | 14 | GLY 13, GLY 15, GLY 60, VAL 14, SER 17, THR 35, GLN 61 |
| Eriodictoyl (S)<sup>Δ*</sup> | *Arachis hypogaea* – fruit, *Artocarpus heterophyllus*- fruit, *Chrysanthemum indicum* –flower, *Citrus* sp., *Cucumis melo* - fruit | IMPHY004038 | -45.517 | 12 | GLY 15, LYS 16, VAL 14, GLY 60, GLY 61, THR 35, GLU 31, ASP 33, SER 17 |
| Plumbagic acid | *Plumbago indica* - root, *Plumbago zeylanica*- whole plant | IMPHY007396 | -45.015 | 11 | VAL 14, GLY 15, SER 17, GLN 61, THR 35 |
| Oleacein | *Jasminum grandiflorum*(Spanish Jasmine)- aerial part | IMPHY013295 | -44.751 | 13 | ASP 33, GLY 60, GLN 61, THR 35, VAL 14, GLY 15, LYS 16, GLY 13 |
| Aristolochic acid C | *Aristolochia tagala* (Indian birthwort) | IMPHY005403 | -44.561 | 14 | GLU 31, ASP 30, VAL 29, VAL 14, GLY 15, GLY 13, SER 17, THR 35 |
| Co-crystal Ligand: Phosphoaminophosphonic acid-guanylate ester (Reference Molecule) | -- | -- | -62.809 | 18 | LYS 16, VAL 29, ASP 30, ASP 119, GLY 13, VAL 14, SER 17, ALA 18, THR 35, ASN 116, LYS 117, ALA 146, GLY 12 |

<sup>#</sup>Common compound; <sup>Δ</sup> enantiomers; \* Selected

### 3.6. Pharmacokinetic and Drug Likeness Screening of Phytochemicals

The drug properties of the top 10 phytochemicals targeting EGFR and HRAS were assessed using the Lipinski Rule of Five and Swiss ADME for drug-likeness (**Table 3**). According to the Lipinski Rule of Five, a compound should satisfy the following criteria: molecular weight (≤ 500 Da), LogP (≤ 5), hydrogen bond donors (≤ 5), hydrogen bond acceptors (≤ 10), and rotatable bonds (≤ 10) (Miebs et al., 2024). Although Molar Refractivity (MR) is not included in these rules, it remains important in drug design, as it reflects a compound’s size, volume, and polarizability, which influence its interaction with targets and membrane permeability (Miebs et al., 2024, Bickerton et al., 2012, Mendie and Hemalatha, 2022). Based on these criteria, all 10 phytochemicals targeting EGFR and HRAS qualify as potential drug candidates (**Table 3**).

**Table 3:**
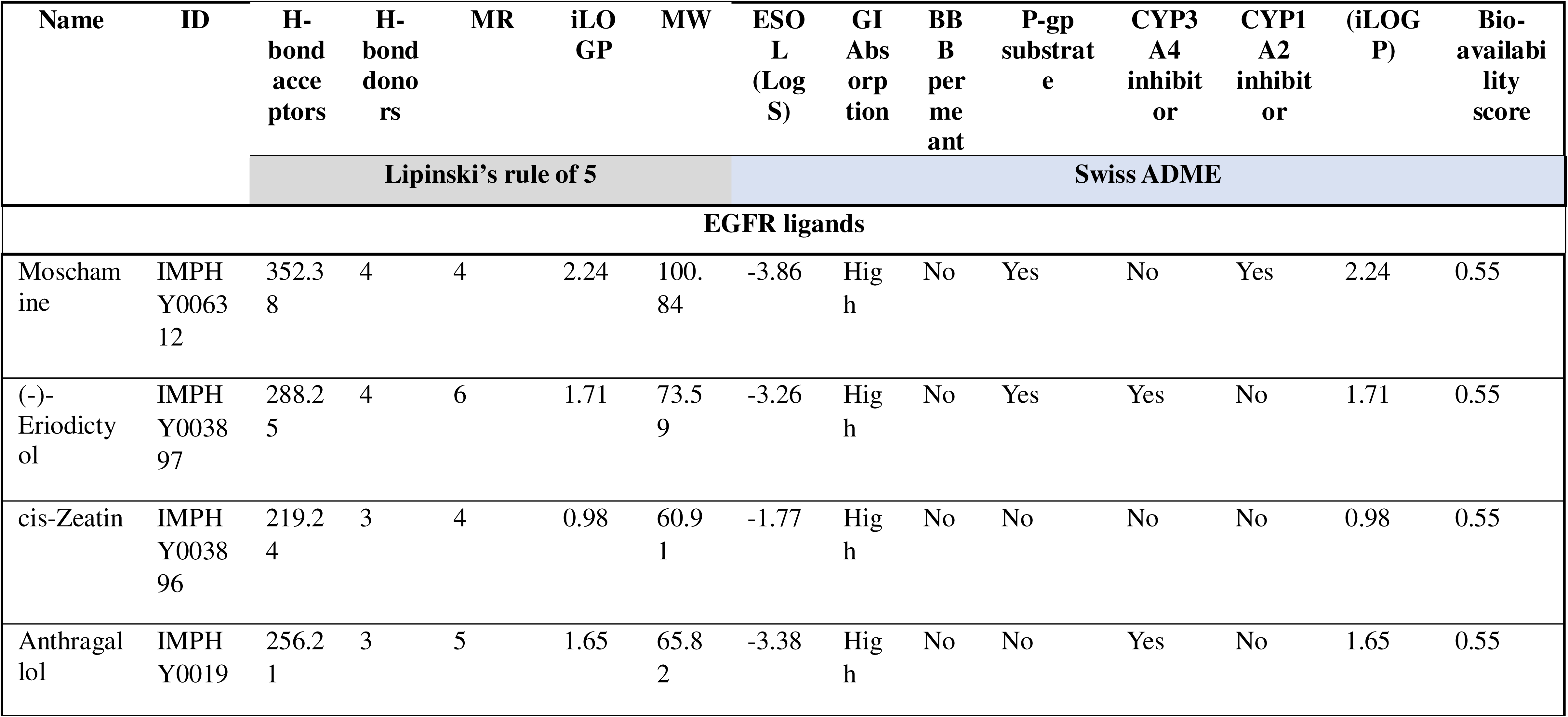
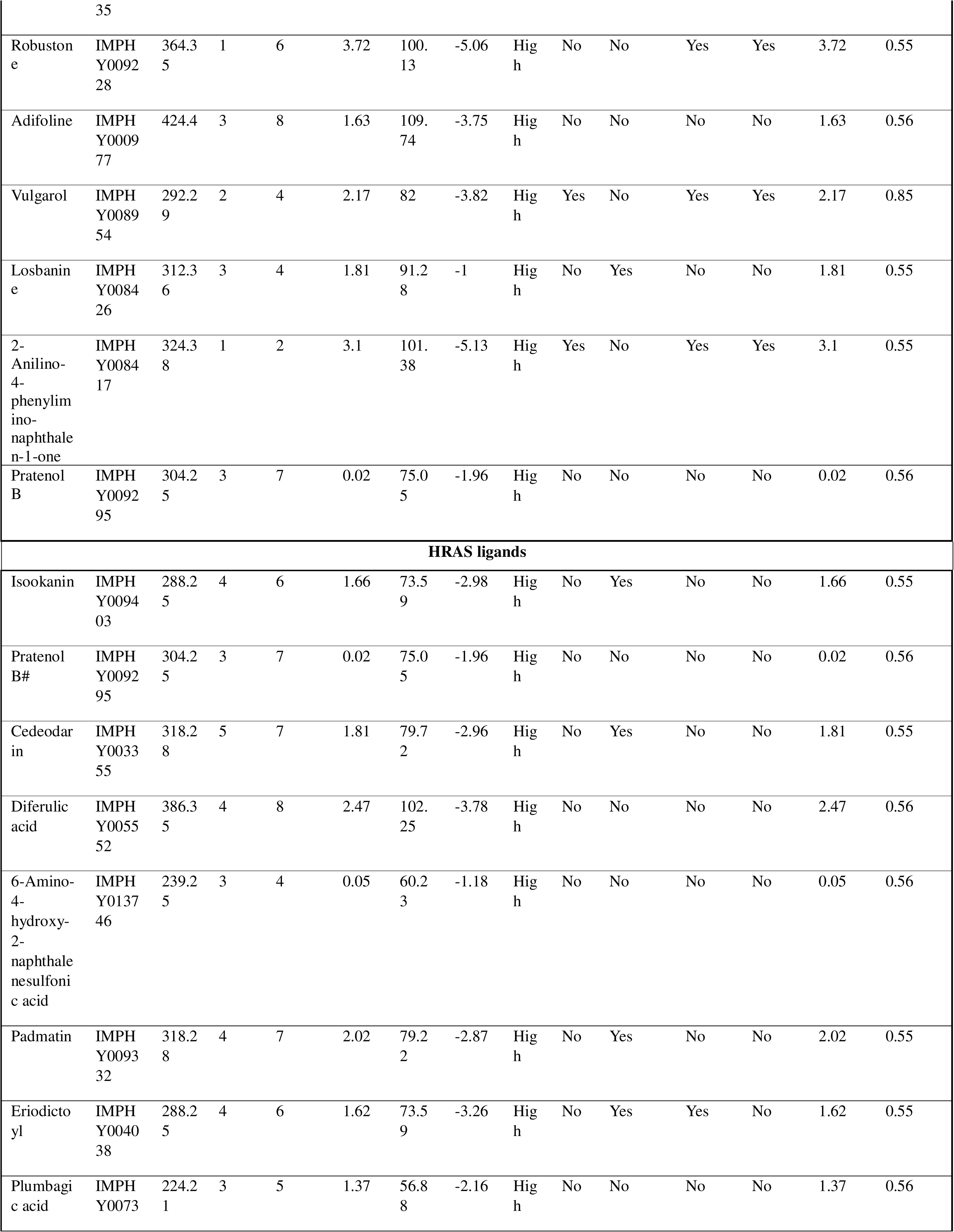
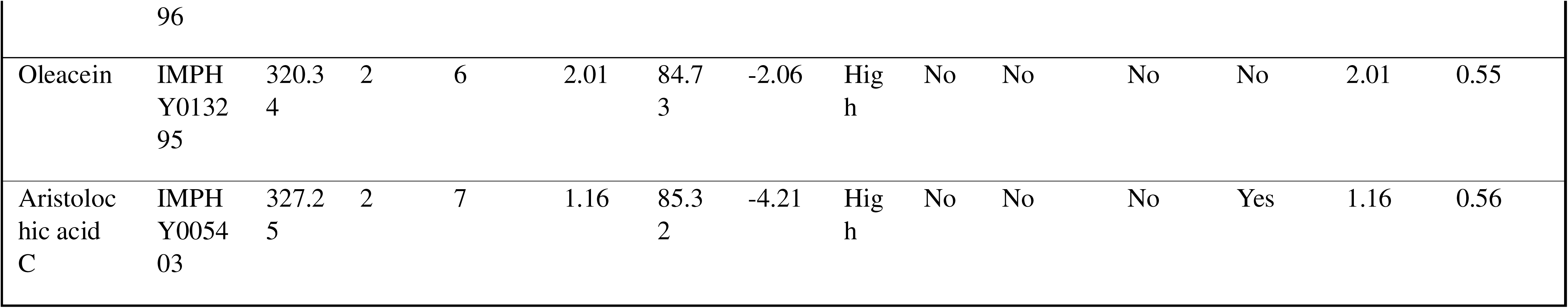
*In silico* pharmacokinetics of EGFR and HRAS ligands

The phytochemicals were screened for ADME (Absorption, Distribution, Metabolism, and Excretion) properties using SwissADME, a free web tool that predicts and evaluates pharmacokinetics and drug-likeness based on multiple computational models. The *in silico* pharmacokinetics of the ligands, as shown in **Table 3**, reveal key insights into their properties.

Most compounds exhibit high to moderate ESOL (Log S) values, indicating good solubility in water (measured in log mol/L). GI absorption values suggest that all compounds are well absorbed in the gastrointestinal (GI) tract, enhancing their oral bioavailability.

The Blood-Brain Barrier (BBB) permeability prediction determines whether a drug can cross the BBB, influencing its potential central nervous system (CNS) activity. Except for Vulgarol and 2-Anilino-4-phenylimino-naphthalen-1-one, all compounds targeting EGFR and HRAS are unable to cross the BBB. This is advantageous, as these drugs are intended to act outside the CNS, reducing the risk of CNS-related side effects.

The P-glycoprotein (P-gp) substrate analysis identifies whether a compound is a P-gp substrate, which affects drug efflux, absorption, and drug resistance. Only a few phytochemicals in the study are P-gp substrates. To overcome P-gp efflux, we suggest potential strategies such as combination therapy with P-gp inhibitors (e.g., Cyclosporine) or utilizing liposomes and lipid nanoparticles (Nguyen et al., 2021).

The CYP3A4 and CYP1A2 inhibition analysis determines whether a compound inhibits cytochrome P450 enzymes (CYP3A4 and CYP1A2), which are essential for drug metabolism and drug-drug interactions. While most compounds are non-inhibitors, a few exhibit inhibitory activity, which can sometimes be beneficial in prolonging drug effects. However, dose adjustments are necessary before administration to avoid potential drug interactions (Lee et al., 2024).

The iLOGP (Intrinsic logP) values indicate that most compounds exhibit optimal lipophilicity, ensuring a good balance between solubility and permeability, which enhances oral absorption.

Finally, the Bioavailability Score, which predicts the probability of oral bioavailability, is greater than 0.5 for all compounds, confirming that they meet the criteria for drug-likeness.

For further verification, we conducted an *in silico* analysis of the pharmacokinetic properties of the top phytochemicals (ligands) identified against EGFR and HRAS using the pkCSM platform. This analysis predicted key pharmacokinetics (PK), toxicity, and ADME (Absorption, Distribution, Metabolism, and Excretion) properties of these small molecules. The pharmacokinetic predictions for the top 10 phytochemicals targeting EGFR and HRAS are presented in **Supplementary Table 1**. Based on these data, we selected the top three phytochemicals for each target: **(-)-Eriodictyol (R), Losbanine, and Pratenol B for EGFR**, and **Pratenol B, Isookanin, and Eriodictoyl (S) for HRAS**. Interestingly, Pratenol B was identified as a common ligand for both targets. The pharmacokinetic properties of these five selected compounds are summarized in **Table 4**.

**Table 4.**
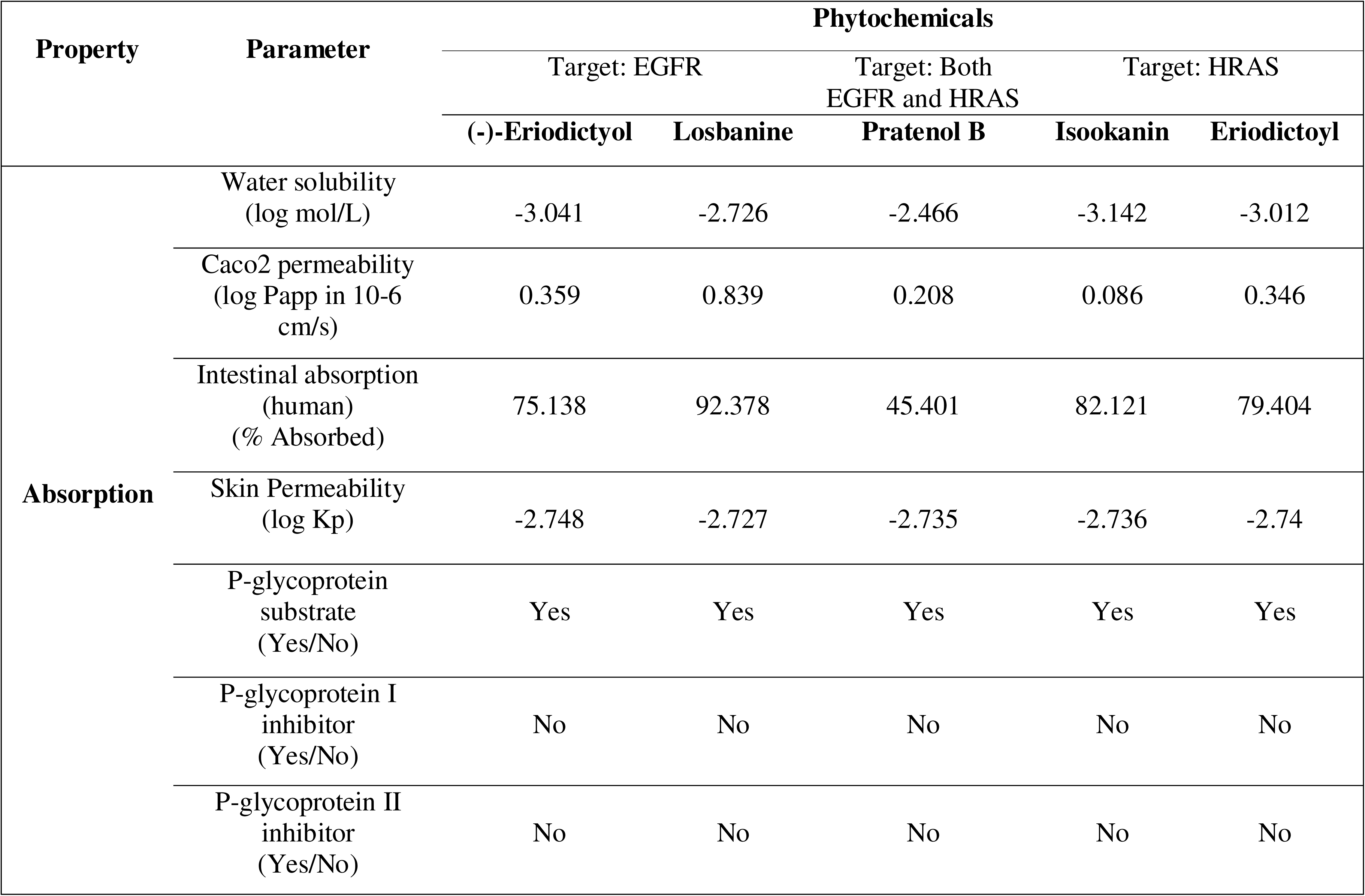
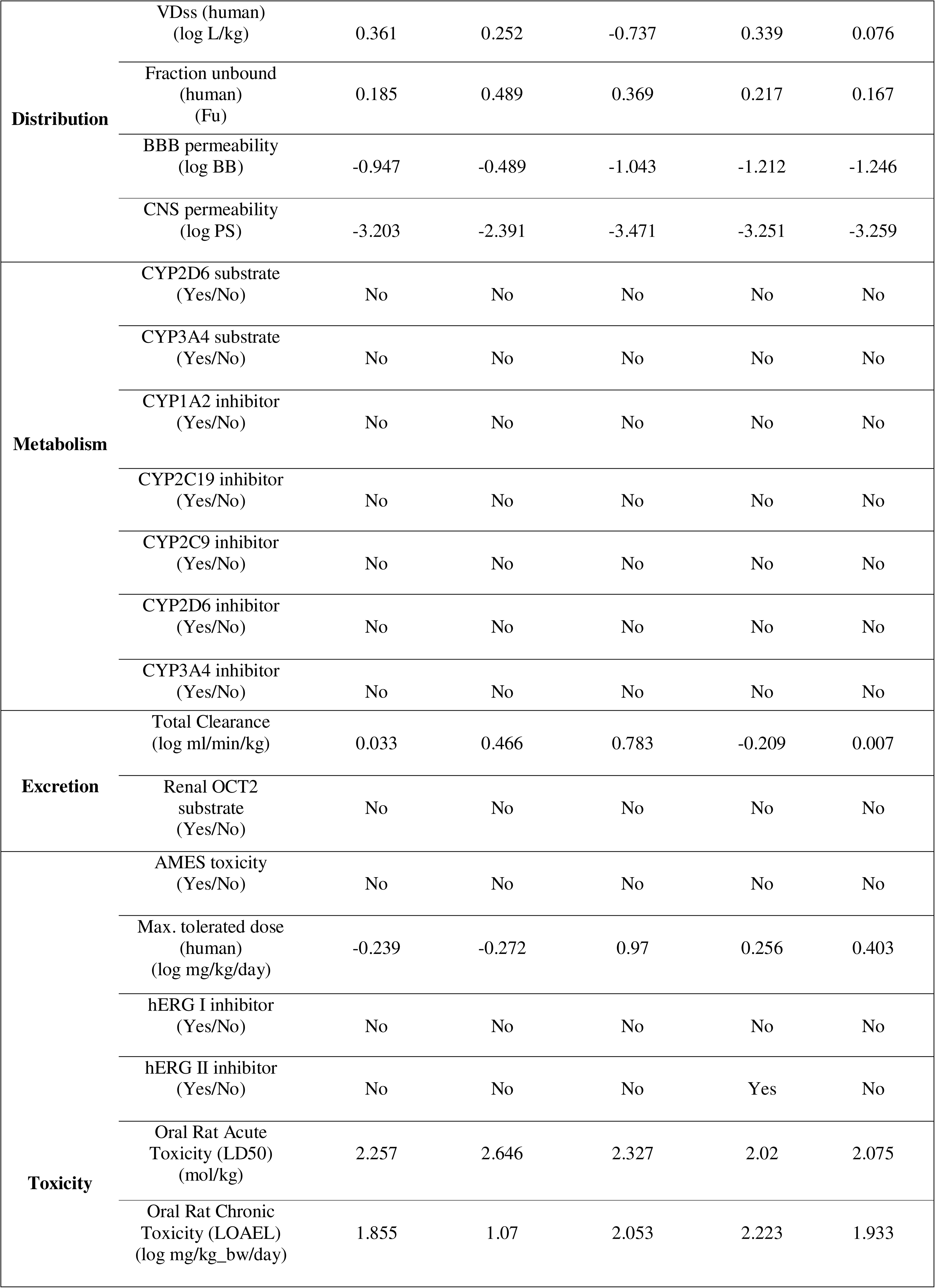
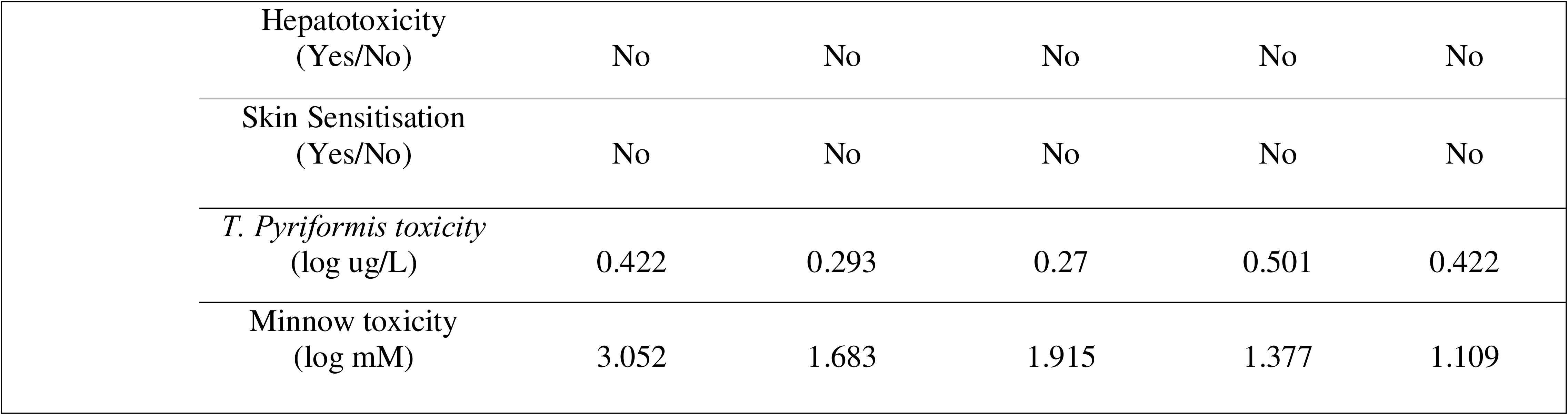
Prediction of pharmacokinetic properties of top phytochemicals (ligands) identified against EGFR and HRAS using pkCSM platform (predicting small-molecule pharmacokinetic properties using graph-based signatures)

The absorption parameters indicate that these compounds exhibit good water solubility, intestinal absorption, and skin permeability, which are essential for effective drug action. Additionally, P-gp efflux, which can hinder drug absorption, can be overcome using various strategies (Nguyen et al., 2021). The VDss (log L/kg) predicts the extent of drug distribution in the body, with moderate VDss being ideal, as extremely high values may lead to tissue accumulation and potential toxicity. All selected compounds are predicted to be BBB and CNS impermeant, reducing concerns related to central nervous system side effects (**Table 4**).

The metabolic profile indicates that none of these drugs are substrates or inhibitors of CYP2D6, CYP3A4, CYP1A2, CYP2C19, or CYP2C9 (**Table 4**). This suggests that they are likely metabolized through alternative pathways such as Flavin-containing Monooxygenases (FMOs), Aldehyde Oxidase (AO), or UDP-glucuronosyltransferase (UGT) (Li et al., 2019). Avoiding CYP-mediated metabolism reduces the risk of drug-drug interactions (DDIs), enhancing their safety when co-administered with other medications (Lee et al., 2024). The excretion parameters suggest that these drugs are not substrates for Organic Cation Transporter 2 (OCT2), indicating that renal excretion is not the primary route of elimination (Wright, 2019). Instead, they are likely cleared via hepatic metabolism or other pathways. Notably, Losbanine and Pratenol B have higher clearance rates, potentially requiring more frequent dosing to maintain therapeutic levels, whereas Isookanin has slow clearance, increasing the risk of drug accumulation, necessitating careful dose adjustments (Lea-Henry et al., 2018).

AMES toxicity predictions suggest that these molecules are non-mutagenic, with a low risk of DNA damage (**Table 4**). The maximum tolerated dose (MTD) (log mg/kg/day) indicates that Pratenol B, Isookanin, and Eriodictoyl have higher max tolerated doses, making them safer and less toxic at therapeutic levels, whereas (-)-Eriodictyol and Losbanine may require careful dose adjustments. The absence of hERG I and II inhibition suggests minimal cardiotoxicity and risk of arrhythmias, although Isookanin may have some toxic effects that warrant careful dosing. LD50 values (>2 mol/kg) indicate moderate-to-low acute toxicity, while LOAEL values (>1 log mg/kg_bw/day) suggest moderate-to-low chronic toxicity. Since all values for hepatotoxicity and skin sensitization are “No,” these molecules are predicted to be safe for the liver and unlikely to cause skin irritation or allergic reactions, making them suitable for dermal applications (**Table 4**).

The *T. Pyriformis* and Minnow toxicity predictions assess the environmental impact of these compounds (Vardhan and Sahoo, 2020). The low toxicity values suggest that these drugs have a minimal ecological risk, making them environmentally safe. Considering the overall pharmacokinetic and toxicity profiles, these phytochemicals exhibit promising drug-likeness properties, although some compounds, such as Isookanin and Losbanine, may require dose adjustments.

### 3.7. Bioactivity Score

The bioactivity score is used to assess the drug-like potential of ligands across various target classes, including G-protein coupled receptors (GPCR), ion channel modulators (ICM), kinase inhibitors (KI), nuclear receptor ligands (NRL), protease inhibitors (PI), and enzyme inhibitors (EI). The Molinspiration online server was employed to predict these scores. The compounds with scores greater than 0.00 exhibit high activity, those ranging between −0.5 and −0.00 indicate moderate activity, while scores lower than −0.5 suggest inactivity (Mendie and Hemalatha, 2022).

**Table 5** presents the bioactivity scores of the top 10 phytochemicals targeting EGFR and HRAS. Among these, the 5 selected compounds demonstrate promising drug-likeness properties, with varied bioactivity across multiple target classes. (-)-Eriodictyol and Eriodictoyl show high bioactivity against NRL and EI, indicating their potential as nuclear receptor modulators with broad pharmacological applications. Losbanine exhibits strong activity for GPCR, ICM, and EI, positioning it as a strong candidate for GPCR-related pathways and enzyme-targeted therapies. Similarly, Isookanin shows high activity in GPCR, NRL, and EI, reinforcing its role in nuclear receptor interactions and enzyme inhibition. Its moderate activity in ICM, KI, and PI suggests a versatile pharmacological profile (**Table 5**).

**Table 5:**
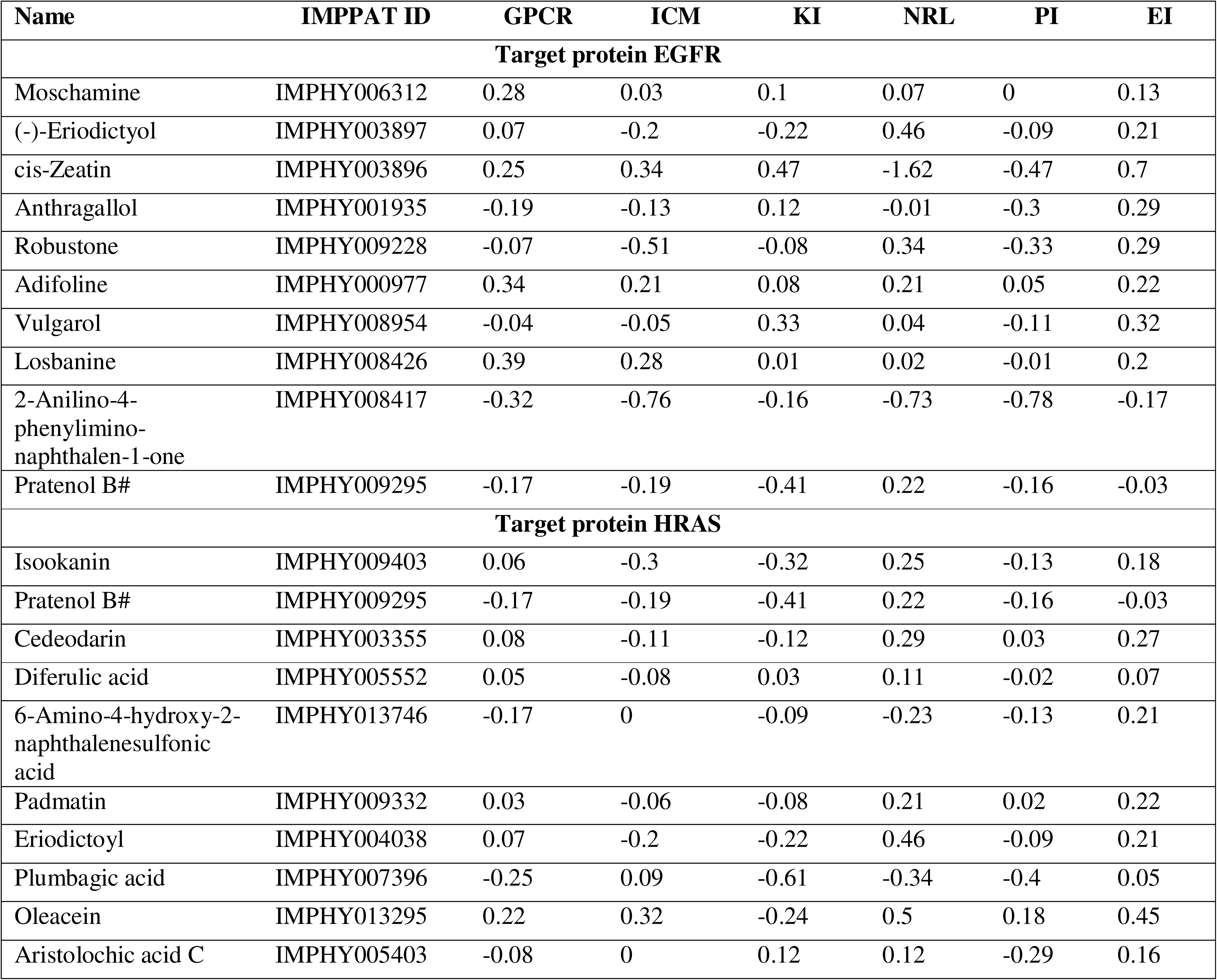
Bioactivity score of compounds targeting EGFR and HRAS

Pratenol B, while moderately active across all target classes, is notable for its broad interaction spectrum, making it a potential multi-target drug candidate (**Table 5**). Its dual activity against EGFR and HRAS further enhances its therapeutic relevance in multi-target treatment strategies.

Overall, these compounds exhibit favorable bioactivity profiles, supporting their potential as promising drug candidates. Among them, Losbanine and Isookanin emerge as highly active compounds, while Pratenol B’s broad interaction potential makes it a noteworthy multi-target agent. However, further experimental validation and optimization will be essential to enhance their drug-like properties and therapeutic efficacy.

### 3.8. Bioavailability Radar

The bioavailability radar is a crucial tool for evaluating the drug-likeness of a compound by analyzing six key physicochemical properties: lipophilicity (LIPO), molecular size (SIZE), polarity (POLAR), insolubility (INSOLU), insaturation (INSATU), and flexibility (FLEX) (Mendie and Hemalatha, 2022, Daina et al., 2017). These parameters collectively determine a compound’s absorption, distribution, metabolism, and excretion (ADME) profile.

Optimal lipophilicity (LIPO) ensures effective membrane permeability, while excessive lipophilicity may lead to toxicity. Molecular size (SIZE) affects absorption, with large molecules struggling to cross membranes, while small ones may be cleared too quickly. Polarity (POLAR) influences solubility and permeability, where highly polar compounds dissolve well but may have poor membrane penetration, whereas low-polarity compounds may suffer from poor solubility. Insolubility (INSOLU) directly impacts absorption, as low solubility can limit bioavailability. Insaturation (INSATU) affects target binding and stability, where higher insaturation enhances biological interactions but may compromise solubility. Flexibility (FLEX) plays a role in binding efficiency, with highly flexible molecules adapting better to targets but potentially losing binding specificity.

A compound’s radar plot should ideally fall within the pink optimal range for good oral bioavailability. Deviations indicate potential pharmacokinetic challenges, necessitating structural modifications to enhance drug-likeness and therapeutic efficacy (Daina et al., 2017, Mendie and Hemalatha, 2022). As shown in **Figure 4**, all selected compounds: (-)-Eriodictyol, Losbanine, and Pratenol B for EGFR, and Pratenol B, Isookanin, and Eriodictoyl for HRAS fall within the optimal range for each parameter. However, a slight increase in INSATU suggests a higher number of double bonds or aromatic rings, which could enhance binding affinity with biological targets without significantly impacting solubility, provided other parameters remain within the optimal range. Given that flexibility (FLEX) and polarity (POLAR) are crucial for bioavailability, and all selected compounds remain within the required limits, they exhibit favorable drug-likeness.

**Figure 4:**
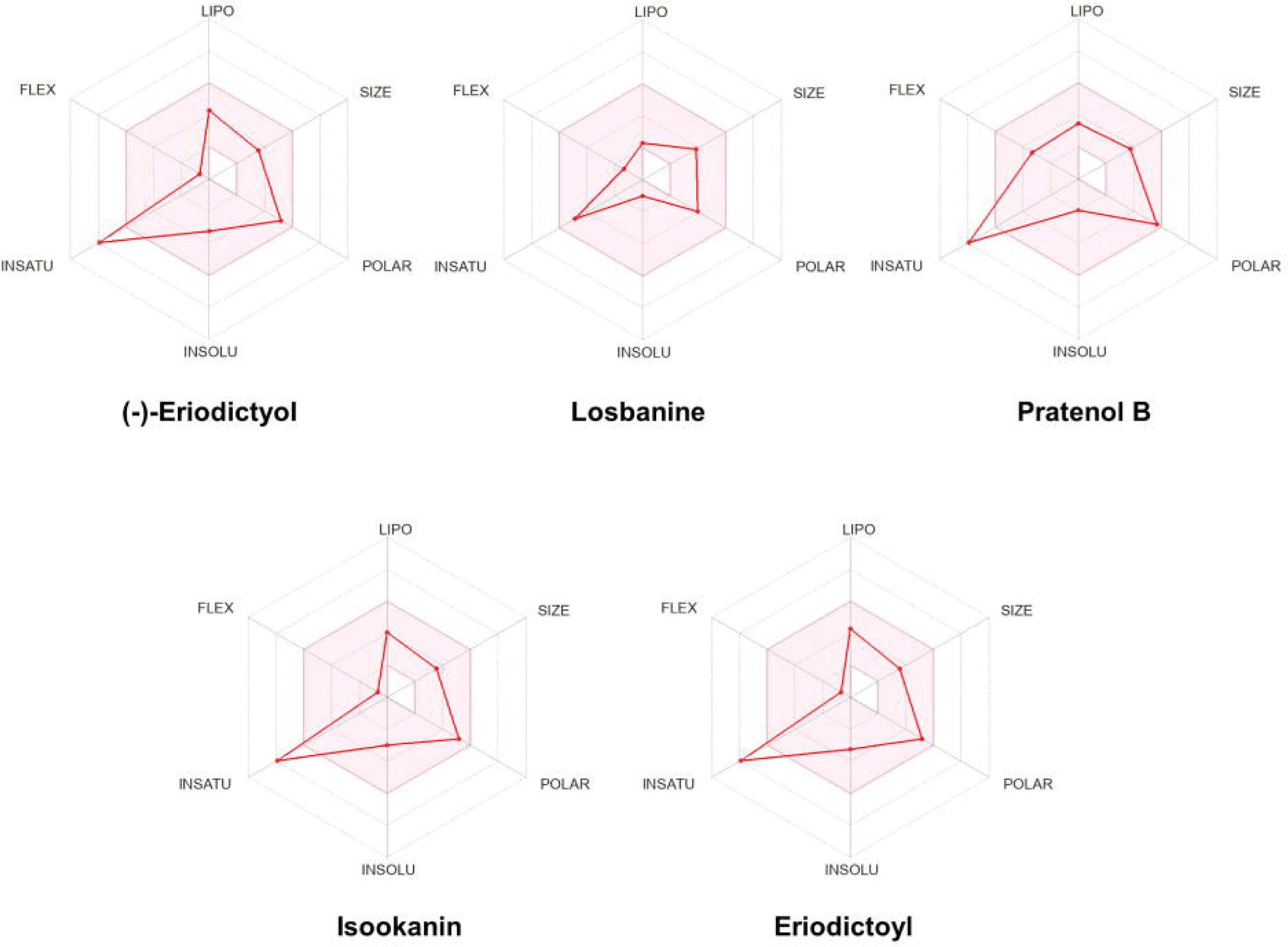
Bioavailability Radar Plot Assessing Drug-Likeness of Selected Compounds for EGFR and HRAS. Bioavailability radar plot illustrating the drug-likeness of selected compounds targeting EGFR [(−)-Eriodictyol (R), Losbanine, and Pratenol B] and HRAS [Pratenol B, Isookanin, and Eriodictyol (S)]. The plot evaluates six key physicochemical properties: lipophilicity (LIPO), molecular size (SIZE), polarity (POLAR), insolubility (INSOLU), insaturation (INSATU), and flexibility (FLEX). The pink-shaded area represents the optimal range for drug-likeness, indicating the suitability of the compounds for oral bioavailability.

### 3.9. Lead Phytochemicals for EGFR and HRAS Inhibition

The selection of (-)-Eriodictyol (R), Losbanine, and Pratenol B for EGFR, and Pratenol B, Isookanin, and Eriodictoyl (S) for HRAS was based on their binding affinity, pharmacokinetics, toxicity, and drug-likeness. Docking simulations from a dataset of 17,000 phytochemicals identified these compounds as top binders, with Pratenol B emerging as a dual inhibitor. (-)-Eriodictyol and Eriodictoyl, being enantiomers, demonstrated high specificity, while Losbanine and Isookanin exhibited favorable molecular interactions. All selected compounds adhered to Lipinski’s Rule of Five and SwissADME criteria, ensuring good solubility, high gastrointestinal (GI) absorption, and minimal blood-brain barrier (BBB) permeability. Their non-inhibition of CYP enzymes reduces the risk of drug-drug interactions (DDIs), while bioavailability radar analysis confirmed optimal drug-likeness, with slight increases in insaturation (INSATU) enhancing target binding. Their 2D and 3D interaction models with their respective targets are presented in **Figure 5**.

**Figure 5:**
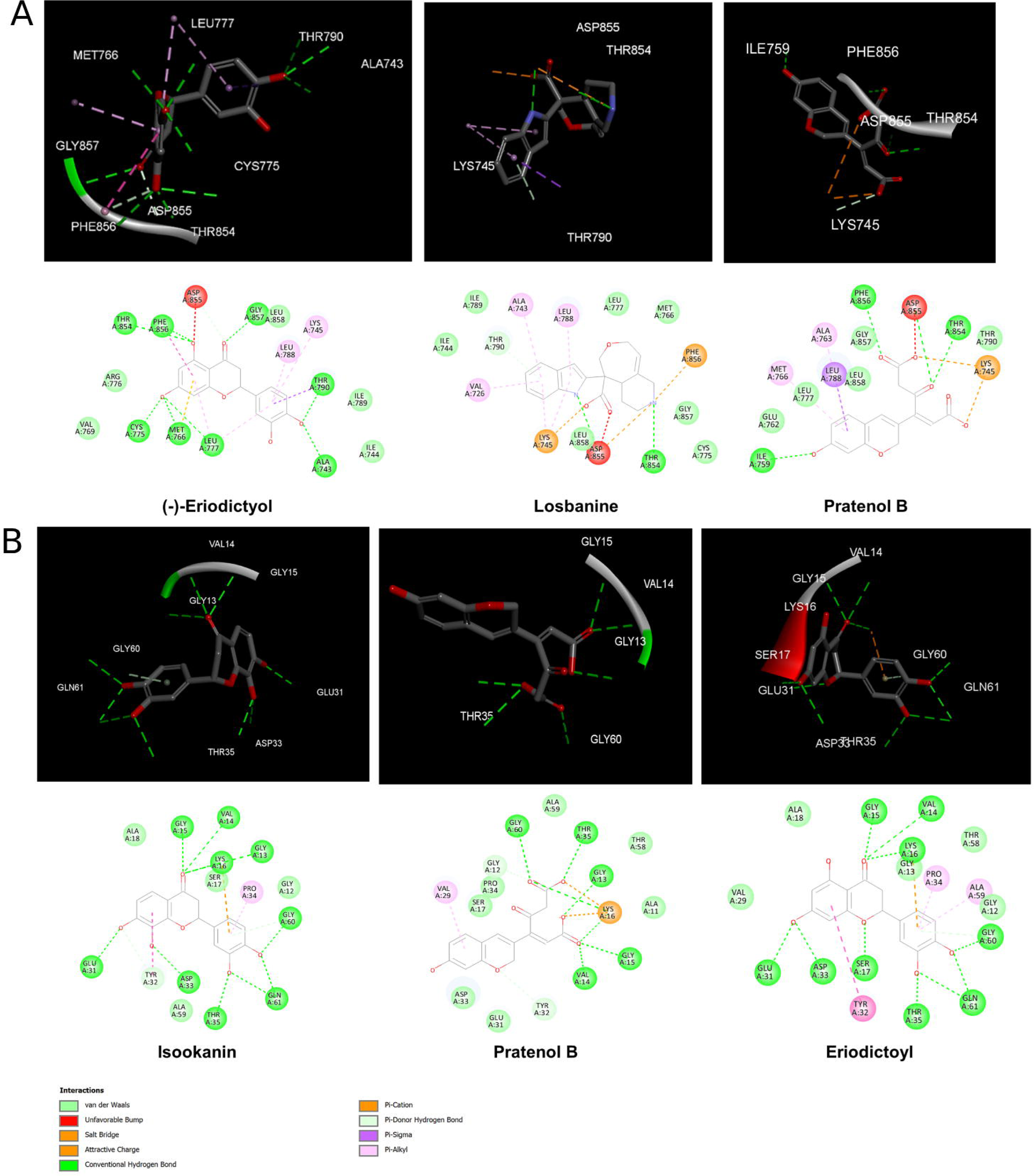
2D and 3D Interaction Models of Selected Compounds with EGFR and HRAS. Molecular docking analysis depicting 2D and 3D interaction models of selected compounds with their respective targets: EGFR [(−)-Eriodictyol (R), Losbanine, and Pratenol B] and HRAS [Pratenol B, Isookanin, and Eriodictyol (S)]. The models illustrate key binding interactions, including hydrogen bonding, hydrophobic interactions, and π-π stacking, providing insights into the molecular recognition and binding affinity of these.

The five selected phytochemicals possess distinct physicochemical properties influencing their drug-likeness, solubility, and bioavailability, as shown in **Supplementary Table 2**. (-)-Eriodictyol, Isookanin, and Eriodictoyl belong to the Flavanone class, while Losbanine is a Carboline alkaloid, and Pratenol B is classified under 1-benzopyrans. Their molecular weights (MW) range from 288.06 Da to 312.15 Da, aligning with Lipinski’s drug-like properties. (-)-Eriodictyol, Isookanin, and Eriodictoyl have six hydrogen acceptors (nHA) and four hydrogen donors (nHD), whereas Losbanine and Pratenol B have fewer hydrogen donors. Pratenol B has the highest number of rotatable bonds, indicating greater molecular flexibility, while (-)-Eriodictyol, Isookanin, and Eriodictoyl (1 rotatable bond) are more rigid.

The Topological Polar Surface Area (TPSA) is highest for Pratenol B (121.13 Å²), reflecting higher polarity, while Losbanine (74.35 Å²) has the lowest TPSA, suggesting enhanced membrane permeability. LogP values, indicating hydrophobicity, range from 0.298 (Losbanine) to 2.373 (-)-Eriodictyol), meaning Losbanine is the most hydrophilic, while (-)-Eriodictyol and Eriodictoyl are more lipophilic. Water solubility (logS) shows that Losbanine (−2.388) and Pratenol B (−2.364) are highly soluble, whereas Eriodictoyl (−3.933) is the least soluble. Acid dissociation constants (pKa Acid) range from 3.338 (Pratenol B) to 8.585 (-)-Eriodictyol), influencing stability at biological pH. Thermal stability varies among the compounds, with Eriodictoyl (264.45°C) exhibiting the highest melting point, while Pratenol B (228.19°C) has the lowest. A similar trend is observed in boiling points, where Eriodictoyl (373.53°C) is the most stable.

Thus, all selected compounds meet key drug-likeness criteria, with Losbanine demonstrating superior solubility, Pratenol B exhibiting higher polarity, and (-)-Eriodictyol, Isookanin, and Eriodictoyl balancing lipophilicity and flexibility. Their favorable solubility, polarity, and bioavailability make them promising drug candidates for further development.

### 3.10. Anticancer activity of Lead Phytochemicals

Eriodictyol (R and S), Losbanine, Isookanin, and Pratenol B have emerged as promising candidates in our study, yet their specific roles in OSCC treatment remain largely unexplored.

Eriodictyol, a naturally occurring flavonoid, is widely present in fruits of medicinal plants (**Table 2**). It exhibits anti-inflammatory, antioxidant, and neuroprotective properties (Li et al., 2020, Yin et al., 2024) and is commercially available, dissolvable in DMSO. Studies have confirmed its anti-tumor activity in lung, colon, breast, pancreatic, liver cancers, and glioma cell lines (Zhang et al., 2020, Li et al., 2020). Eriodictyol inhibits glioma cell proliferation, migration, and invasion while inducing apoptosis by suppressing the PI3K/Akt/NF-κB signaling pathway, leading to the downregulation of p-PI3K, p-Akt, p-IκBα, and p-NF-κB (Li et al., 2020). Its anti-glioma effects were further validated in a xenograft mouse model, where it effectively suppressed tumor growth by reducing proliferation and promoting apoptosis (Li et al., 2020). Recent findings highlight Eriodictyol’s potential in OSCC, as it has been reported to inhibit the proliferation and induce apoptosis in SCC131 cell line via the PI3K/Akt pathway (Xu et al., 2024). However, no detailed mechanistic or therapeutic studies have been conducted against OSCC, presenting an opportunity for further investigation.

Losbanine and Isookanin remain relatively novel, with no reported anticancer activity, warranting further exploration of their therapeutic potential. Pratenol B, derived from *Trifolium pratense* (red clover), is also largely unexplored for cancer therapy. However, *Trifolium pratense* extract has demonstrated significant anticancer properties, particularly in NALM-6 cells (B-cell acute lymphoblastic leukemia), where it induces autophagy and apoptosis pathways, suggesting its potential for ALL treatment (Shirani Asl et al., 2024).

Given their novelty in OSCC research, further studies are crucial to assess their therapeutic mechanisms and clinical relevance, positioning them as promising candidates for future cancer treatment strategies.

### 3.11. Molecular Dynamics Simulation

To further assess the stability of the protein-ligand complexes following evaluation of their drug-like properties, molecular dynamics (MD) simulations were performed. Simulations were conducted for EGFR in complex with (-)-Eriodictyol (R), Losbanine, Pratenol B, and the reference molecule Gefitinib (**Figure 6A-D** and **7A-D**). The results revealed that both Pratenol B and Gefitinib formed stable complexes with EGFR throughout the 100 ns simulation. The EGFR–Pratenol B complex achieved stability around 10 ns and maintained consistent interactions with the receptor throughout the simulation period, indicating a potentially long-lasting binding affinity (**Figure 6C**). Gefitinib also displayed stable binding, although it exhibited slightly more fluctuations, suggesting minor deviations from the active site during the simulation (**Figure 6A**).

**Figure 6.**
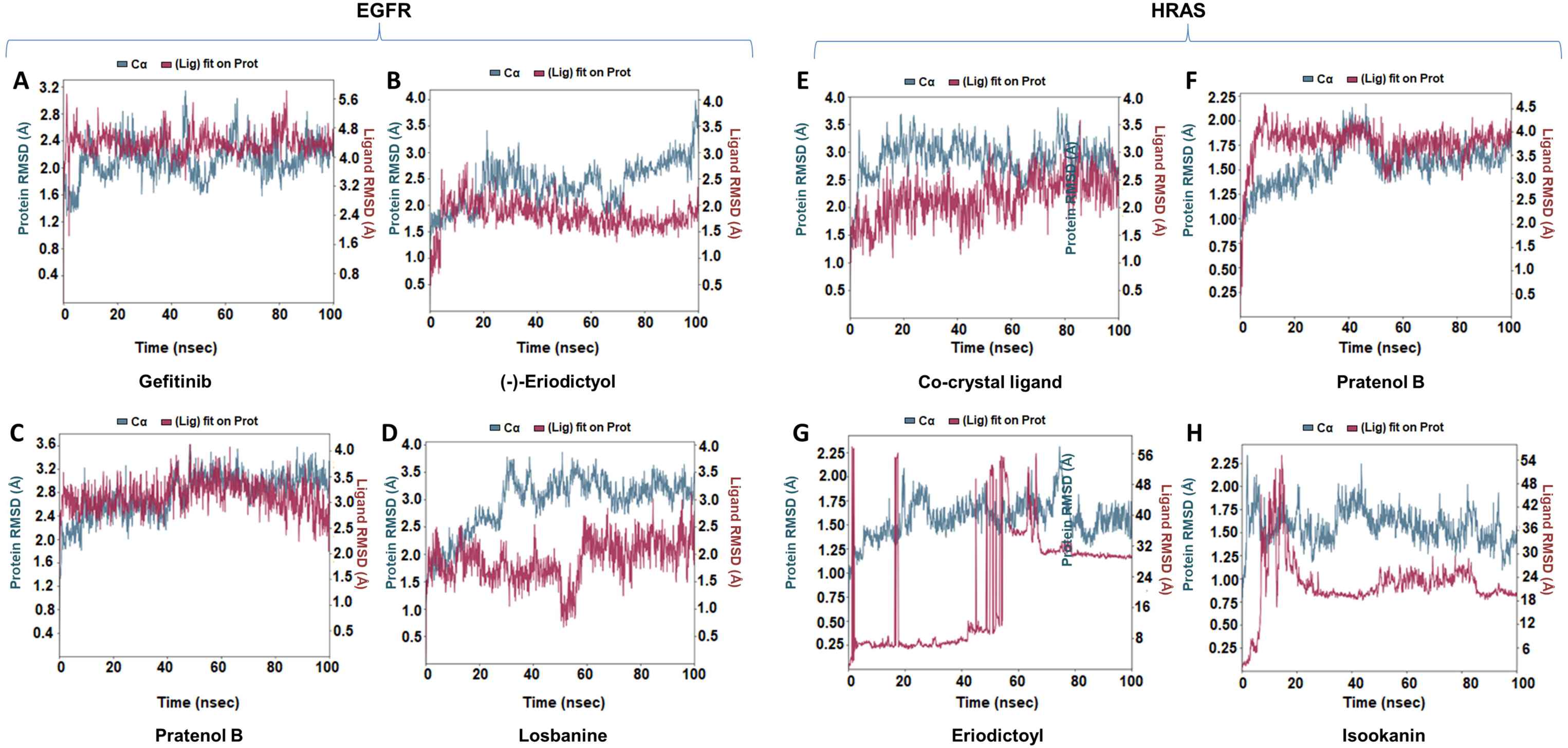
RMSD analysis of protein–ligand complexes for EGFR and HRAS over 100 ns molecular dynamics simulations. **(A–D)** shows the Root Mean Square Deviation (RMSD) profiles for EGFR complexes with different ligands, and (**E–H**) represents HRAS complexes. In each panel, the blue line represents the protein backbone RMSD (Cα atoms), while the maroon line represents the ligand RMSD when fitted onto the protein. The stability of each complex is inferred from the RMSD profiles over the 100 ns simulation.

In contrast, (-)-Eriodictyol (R) remained stably bound within the active site up to approximately 55 ns, after which it began to drift away and was completely dislodged from the active site by the end of the simulation (**Figure 6B**). Losbanine demonstrated substantial fluctuations starting from 20 ns, indicating poor stability and weak retention within the binding pocket (**Figure 6D**).

The RMSF analysis for all EGFR-ligand complexes revealed fluctuations across individual residues (scale: 0–700 residue positions) over the 100 ns simulation (**Figure 7A-D**). Notably, the highest fluctuations were observed at the active site residues. Asp855, which interacts with all the ligands, showed the most prominent fluctuation. In the Pratenol B complex, Met766 exhibited greater flexibility compared to its behavior in the Eriodictyol (R) and Losbanine complexes, suggesting that bond flexibility may contribute to enhanced complex stability. These findings are consistent with the docking results, which also indicated significant interactions at these residues.

**Figure 7.**
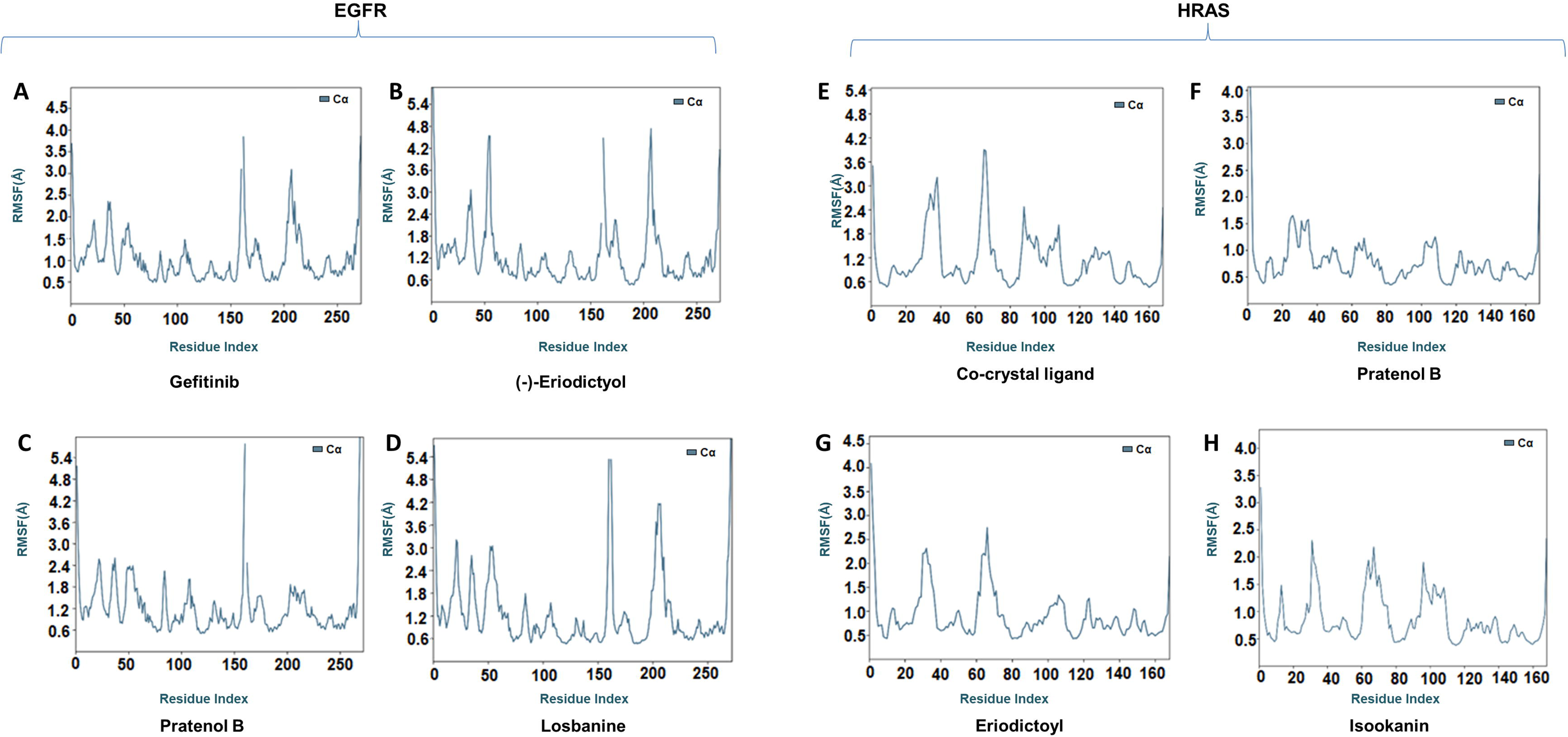
RMSF analysis of protein–ligand complexes for EGFR and HRAS over 100 ns molecular dynamics simulations. (**A–D**) display the Root Mean Square Fluctuation (RMSF) of Cα atoms in EGFR upon binding with different ligands, and (**E–H**) shows RMSF plots for HRAS in complex with different ligands. RMSF values are plotted against the residue index, indicating the flexibility of individual amino acid residues during the simulation.

Similar MD simulations were conducted for HRAS in complex with Pratenol B, Isookanin, Eriodictyol (S), and the co-crystal ligand Phosphoaminophosphonic acid-guanylate ester as a reference (**Figure 6E-H** and **7E-H**). The simulation revealed that Isookanin and Eriodictyol (S) were highly unstable and showed significant fluctuations throughout the simulation period. Initially, the HRAS–Pratenol B complex exhibited fluctuations in both the ligand and receptor; however, the complex reached stability after 37 ns and remained stable for the remainder of the 100 ns run (**Figure 6F**). The reference ligand also showed initial instability until around 62 ns, after which it stabilized, suggesting a conformational adjustment period before achieving a stable binding state (**Figure 6E**).

RMSF analysis of the HRAS complexes revealed that residues in the 20–40 range were more flexible across all ligand-bound structures, while residues in the 60–80 range exhibited reduced fluctuation in the Pratenol B complex compared to the reference ligand **(Figure 7E-H)**.

Overall, the MD simulation studies demonstrated that both EGFR and HRAS formed the most stable complexes with **Pratenol B**, outperforming the reference compounds in terms of binding stability and interaction consistency.

## 4. Conclusion

This study presents a transcriptomic analysis of OSCC in Indian patients, highlighting MAPK pathway dysregulation as a key driver of tumor progression. Within this pathway, EGFR and HRAS emerged as top crucial therapeutic targets; however, existing treatments are limited by drug resistance and toxicity. Given the scarcity of HRAS inhibitors and the challenges associated with EGFR-targeting therapies, there is a pressing need for novel small-molecule inhibitors with fewer side effects.

To address this, a natural product-based drug discovery approach was employed through docking simulations, screening 17,000 phytochemicals. This led to the identification of (-)-Eriodictyol (R), Losbanine, and Pratenol B for EGFR, and Pratenol B, Isookanin, and Eriodictoyl (S) for HRAS inhibition. Pratenol B demonstrated dual inhibition of both targets and most stable complex, while Eriodictyol (R and S) exhibited strong binding affinity and favorable drug-likeness. All selected compounds displayed optimal pharmacokinetics, high bioavailability, and low toxicity, ensuring both safety and efficacy.

Harnessing India’s rich medicinal plant resources could facilitate the development of cost-effective, plant-derived inhibitors for OSCC treatment. Future research should focus on experimental validation in *in vitro* and *in vivo* OSCC models, optimization of bioavailability and formulation strategies, and combination therapy approaches to enhance treatment efficacy and overcome drug resistance. Clinical translation and safety evaluations will be essential to advance these compounds as potential therapeutic agents. With continued research and development, these phytochemicals hold significant promise for delivering safer, more effective, and accessible treatments for OSCC and other malignancies.

## Supplementary Materials

**Supplementary Table 1: Prediction of pharmacokinetic properties of phytochemicals (ligands) identified against EGFR and HRAS using pkCSM platform**

**Supplementary Table 2: Physicochemical properties of selected phytochemicals**

**Supplementary Figure 1: Key quality control metrics for RNA sequencing, and summary of transcriptomic alterations.**

**Supplementary Figure 2: Chemical structures of the top 10 selected phytochemicals targeting EGFR and HRAS.**

## Author Contributions

Conceptualization: S. Sur and D. Davray; Data Analysis and Interpretation: S. Behara, U. Yadav, S. Nagar, V. K. Sahu, S. Basu, D. Davray, and S. Sur; Sample Procurement: S. Gupta and B. M. Rudagi; Histopathological Evaluation: S. Kheur; Writing – Original Draft Preparation: S. Behara, U. Yadav, S. Nagar, V. K. Sahu, S. Sur, and D. Davray. All authors have read and approved the final version of the manuscript.

## Data Availability

All data generated in this study are included within the manuscript.

## Supporting information

Supplementary Figure 1: Key quality control metrics for RNA sequencing, and summary of transcriptomic alterations.

Supplementary Figure 2: Chemical structures of the top 10 selected phytochemicals targeting EGFR and HRAS.

Supplementary Table 1: Prediction of pharmacokinetic properties of phytochemicals (ligands) identified against EGFR and HRAS using pkCSM platform

Supplementary Table 2: Physicochemical properties of selected phytochemical

## Acknowledgment

We sincerely thank the Director of Dr. D. Y. Patil Biotechnology and Bioinformatics Institute, Dr. D. Y. Patil Vidyapeeth, Tathawade, Pune, for invaluable support and encouragement. We also acknowledge the Department of Science & Technology (DST), Government of India, for providing the DST-FIST grant (SR/FST/LS-I/2017/70), which facilitated research infrastructure and instrumentation.

## Declarations

### Ethics Approval

This study was approved by the Institutional Biosafety Committee (IBSC) [DYPBBI/1/2023 dated 7/1/2023] and Ethics Committee [DYPV/EC/910/23 dated 23/ January/ 2023], Dr. D.Y. Patil Biotechnology and Bioinformatics Institute, Dr. D. Y. Patil Vidyapeeth (DPU), Pune, India.

## Abbreviations

ADMET: Absorption, Distribution, Metabolism, Excretion, and Toxicity
AMPK: AMP-Activated Protein Kinase
AREG: Amphiregulin
BBB: Blood-Brain Barrier
CDC25B: Cell Division Cycle 25B
c-Myc: Cellular Myelocytomatosis Oncogene
CNS: Central Nervous System
DEGs: Differentially Expressed Gene
DMSO: Dimethyl Sulfoxide
EGFR: Epidermal Growth Factor Receptor
FDA: Food and Drug Administration
GO: Gene Ontology
GCO: Global Cancer Observatory
HRAS: Harvey Rat Sarcoma Viral Oncogene Homolog
HSPA8: Heat Shock Protein Family A (Hsp70) Member 8
hERG I/II: Human Ether-à-go-go-Related Gene Potassium Channel Inhibitors (hERG I and II)
IMPPAT: Indian Medicinal Plants, Phytochemistry, and Therapeutics
IL1A: Interleukin 1 Alpha
KRAS: Kirsten Rat Sarcoma Viral Oncogene Homolog
KEGG: Kyoto Encyclopedia of Genes and Genomes
MET: MET Proto-Oncogene, Receptor Tyrosine Kinase
MAPK: Mitogen-Activated Protein Kinase
NRAS: Neuroblastoma Rat Sarcoma Viral Oncogene Homolog
NF-κB: Nuclear Factor Kappa B
OSCC: Oral squamous cell carcinoma
PRAD-1 (also known as Cyclin D1): Parathyroid Adenoma 1
P-gp: P-Glycoprotein
pkCSM: pharmacokinetics-chemical space modeling
PI3K: Phosphoinositide 3-Kinase
PDB: Protein Data Bank
RMSD: Root Mean Square Deviation
RMSF: Root Mean Square Fluctuation
TGFA: Transforming Growth Factor Alpha
VEGFC: Vascular Endothelial Growth Factor C

## Supplementary Figures

**Supplementary Figure 1: (A–B)** Key quality control metrics for RNA sequencing, including filtered reads and alignment summary. **(C)** Summary of transcriptomic alterations in OSCC tumor samples (T) compared to adjacent non-tumor tissues (N), highlighting differentially expressed genes.

**Supplementary Figure 2:** Chemical structures of the top 10 selected phytochemicals targeting EGFR and HRAS, visualized using JSmol retrieved from the IMPPAT database (https://cb.imsc.res.in/imppat/).

