## Supplementary figures and images for "Integrative Transcriptomics and Phytochemical Screening Reveal Pratenol B, Eriodictyol, Losbanine, and Isookanin, as Potential EGFR and HRAS Inhibitors in Indian Oral Squamous Cell Carcinoma Patients"

### Supplementary Figure 1: Key quality control metrics for RNA sequencing, and summary of transcriptomic alterations.

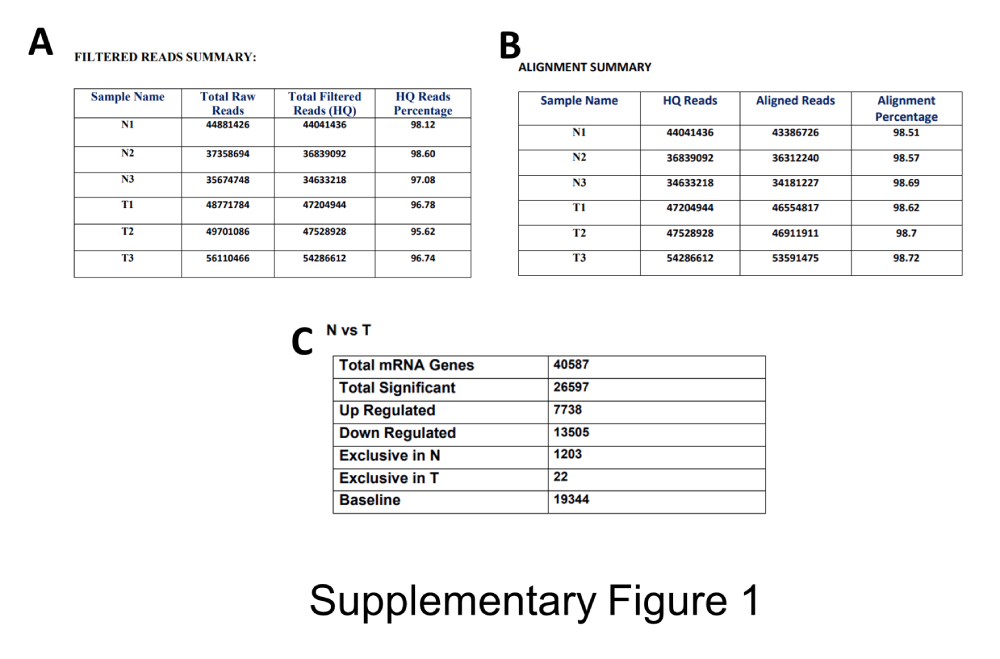

### Supplementary Figure 2: Chemical structures of the top 10 selected phytochemicals targeting EGFR and HRAS.

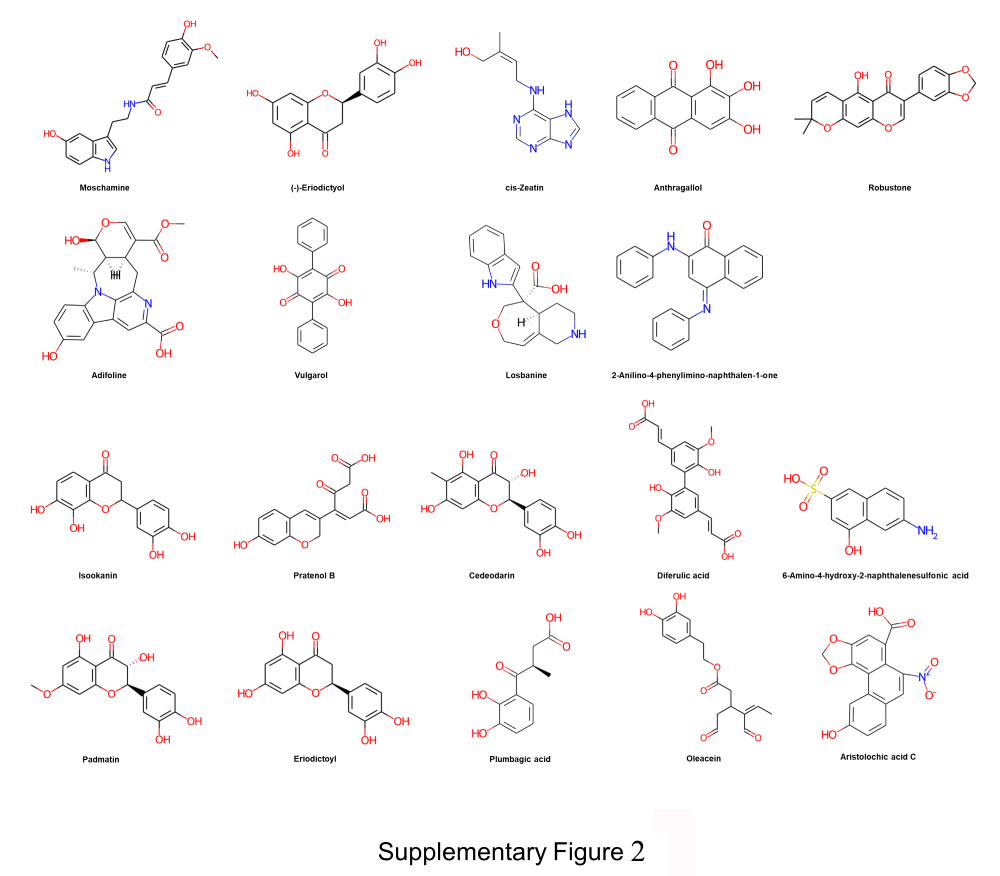
