## Supplementary Table 2: Physicochemical properties of selected phytochemical for "Integrative Transcriptomics and Phytochemical Screening Reveal Pratenol B, Eriodictyol, Losbanine, and Isookanin, as Potential EGFR and HRAS Inhibitors in Indian Oral Squamous Cell Carcinoma Patients"

**Supplementary Table 2**: Physicochemical properties of selected phytochemicals

| **Property** | **(-)-Eriodictyol** | **Losbanine** | **Pratenol B** | **Isookanin** | **Eriodictoyl** |
| --- | --- | --- | --- | --- | --- |
| Chemical  Class | Flavanones | Carboline alkaloids | 1-benzopyrans | Flavanones | Flavanones |
| MW | 288.06 | 312.15 | 304.06 | 288.06 | 288.06 |
| nHA | 6 | 5 | 7 | 6 | 6 |
| nHD | 4 | 3 | 3 | 4 | 4 |
| nRot | 1 | 2 | 5 | 1 | 1 |
| nRing | 3 | 4 | 2 | 3 | 3 |
| MaxRing | 10 | 11 | 10 | 10 | 10 |
| nHet | 6 | 5 | 7 | 6 | 6 |
| fChar | 0 | 0 | 0 | 0 | 0 |
| nRig | 18 | 23 | 15 | 18 | 18 |
| Flexibility | 0.056 | 0.087 | 0.333 | 0.056 | 0.056 |
| Stereo Centers | 1 | 2 | 0 | 1 | 1 |
| TPSA | 107.22 | 74.35 | 121.13 | 107.22 | 107.22 |
| logS | -3.824 | -2.388 | -2.364 | -3.743 | -3.933 |
| logP | 2.373 | 0.298 | 0.451 | 1.914 | 2.205 |
| logD7.4 | 2.44 | 1.158 | 0.876 | 2.059 | 2.273 |
| pka (Acid) | 8.585 | 6.962 | 3.338 | 7.807 | 8.356 |
| pka (Base) | 5.687 | 6.945 | 4.018 | 3.076 | 4.346 |
| Melting point | 245.905 | 234.532 | 228.193 | 247.558 | 264.45 |
| Boiling point | 350.172 | 304.985 | 303.048 | 354.559 | 373.533 |

Molecular weight (MW), number of hydrogen acceptors (nHA), number of hydrogen donors (nHD), number of rotatable bonds (nRot), number of ring systems (nRing), size of the largest ring in the compound (MaxRing), number of heteroatoms (nHet), formal charge of the molecule (fChar), number of rigid bonds in structure (nRig), topological polar surface area (TPSA), logarithm of water solubility (logS), partition coefficient bet. n-octanol and water (hydrophobicity), distribution coefficient at pH 7.4 (logD7.4), acid dissociation constant (pka (Acid)), base dissociation constant (pka (Base))
